# Dispersal inference from population genetic variation using a convolutional neural network

**DOI:** 10.1101/2022.08.25.505329

**Authors:** Chris C. R. Smith, Silas Tittes, Peter L. Ralph, Andrew D. Kern

**Author notes:** April 11, 2023.

## Abstract

The geographic nature of biological dispersal shapes patterns of genetic variation over landscapes, making it possible to infer properties of dispersal from genetic variation data. Here we present an inference tool that uses geographically distributed genotype data in combination with a convolutional neural network to estimate a critical population parameter: the mean per-generation dispersal distance. Using extensive simulation, we show that our deep learning approach is competitive with or outperforms state-of-the-art methods, particularly at small sample sizes. In addition, we evaluate varying nuisance parameters during training—including population density, demographic history, habitat size, and sampling area—and show that this strategy is effective for estimating dispersal distance when other model parameters are unknown. Whereas competing methods depend on information about local population density or accurate identification of identity-by-descent tracts, our method uses only single-nucleotide-polymorphism data and the spatial scale of sampling as input. Strikingly, and unlike other methods, our method does not use the geographic coordinates of the genotyped individuals. These features make our method, which we call “`disperseNN`”, a potentially valuable new tool for estimating dispersal distance in non-model systems with whole genome data or reduced representation data. We apply `disperseNN` to 12 different species with publicly available data, yielding reasonable estimates for most species. Importantly, our method estimated consistently larger dispersal distances than mark-recapture calculations in the same species, which may be due to the limited geographic sampling area covered by some mark-recapture studies. Thus genetic tools like ours complement direct methods for improving our understanding of dispersal.

## Introduction

Organisms vary greatly in their capacity to disperse across geographic space. Indeed, the movement of individuals or of gametes across a landscape helps to determine the spatial scale of genetic differentiation and the spread of adaptive variants across natural populations (Broquet and Petit, 2009). Consequently, understanding dispersal is relevant for conservation biology (Driscoll et al., 2014), studying climate change response and adaptation (Travis et al., 2013), managing invasive and disease vector populations (Harris et al., 2009; Orsborne et al., 2019), phylogeography (Kadereit et al., 2005), studying hybrid zones and microbial community ecology (Barton, 1979; Evans et al., 2017), and for parameterizing models in ecology and evolution (Barton et al., 2002). Despite the importance of dispersal, it remains challenging to obtain estimates for dispersal distance in many species.

Some methods infer dispersal distance by directly observing individual movement, using radio-tracking technology, or by tagging and recapturing individuals in the field. However, such measurements can be expensive to obtain and lead to estimates with high uncertainty. Furthermore, they do not always provide a complete picture of *effective* dispersal rate—that is, how far successfully-reproducing individuals travel from their birth location (for a review, see Bradburd and Ralph, 2019). Effective dispersal is relevant for describing the movement of genetic material across geography, which can be important for understanding population structure, connectivity between populations, evolutionary dynamics of selected alleles, and changes to a species’ range (Slatkin, 1987; Peacock, 1997).

Another type of method infers effective dispersal distance from genotypes of a single temporal sample, without directly observing movement of individuals. Such inference is possible because population genetics theory predicts how demographic parameters such as the rate of gene flow across the landscape affect the genetic variation of a population (Barton et al., 2013). To infer dispersal distance, current population-genetics­based estimators (Rousset, 1997; Ringbauer et al., 2017) use geographically referenced DNA sequences and can obtain useful estimates of the per-generation dispersal distance, without the need for tracking or recapturing individuals.

Importantly, current population-genetics-based estimators require additional data that can be prohibitively expensive, especially for non-model species: for example, an independent estimate of population density (Rousset, 1997), or genomic identity-by-descent blocks (Ringbauer et al., 2017). Specifically, the seminal method of Rousset (1997) is designed for estimating neighborhood size, *N_loc_*, which can be thought of as the number of neighboring individuals or potential mates that are within a few multiples of the dispersal distance (Wright, 1946). Wright defined neighborhood size as *N_loc_ =* 4*πDσ*^2^, where *σ* is the dispersal distance and *D* is the population density. Therefore the accuracy of Rousseťs method depends on having a good *a priori* estimate of population density. One way to jointly infer dispersal and density works by modeling genomic identity-by-descent tracts (e.g., Barton et al., 2013; Baharian et al., 2016; Ringbauer et al., 2017). The program MAPS (Al-Asadi et al., 2019) uses identity-by-descent information to infer heterogeneous dispersal and density across a landscape. Similarly, inferred tree sequences can now be used to infer dispersal rate (Osmond and Coop, 2021). Although powerful when applied to high quality data, the latter methods are limited by the availability of confident identity-by-descent blocks and tree sequences; these data types remain unavailable or difficult to estimate accurately for most species.

Another type of population-genetics-based method estimates *relative* migration rates, for example EEMS (Petkova et al., 2016), FEEMS (Marcus et al., 2021), and other landscape genetics tools. Although such methods work well for some applications, such as identifying barriers to dispersal, they don’t inform us about the magnitude of dispersal (and so results are not returned with units such as meters per generation). Furthermore, these and related tools model gene flow using an approximate analogy to electrical resistance which can produce misleading results especially in the presence of biased migration (Lundgren and Ralph, 2019). Yet another class of dispersal estimation uses non-recombining DNA segments (Neigel et al., 1991; Neigel and Avise, 1993; Lemey et al., 2010), which is valuable for studying phylogeography of viruses, or studying the movement of mitochondrial DNA or sex chromosomes. However, in recombining species we would ideally leverage more than one locus to infer dispersal rate. In the current paper we set out to develop a method for estimating dispersal distance that can be applied widely, including in non-model species without good assemblies or knowledge of population density.

To do this we use simulation-based inference via deep learning to infer dispersal from genotype data. Deep learning is a form of supervised machine learning that builds a complex function between input and output involving successive layers of transformations through a “deep” neural network. An important advantage of this class of methods is their ability to handle many correlated input variables without knowledge of the variables’ joint probability distribution. Like all supervised machine learning methods, deep neural networks can be trained on simulated data, which bypasses the need to obtain empirical data for training (Schrider and Kern, 2018). Over the past few years, deep learning has been used in a number of contexts in population genetics: for example, inferring demographic history in *Drosophila* (Sheehan and Song, 2016), detection of selective sweeps (Kern and Schrider, 2018), detecting adaptive introgression in humans (Gower et al., 2021), identifying geographic origin of an individual using their DNA (Battey et al., 2020a), and estimating other population genetic parameters like recombination rate (Flagel et al., 2019; Adrion et al., 2020).

We present the first use of deep learning for estimation of spatial population genetic parameters. Our method, called `disperseNN`, uses forward in time spatial genetic simulations (Haller and Messer, 2019; Battey et al., 2020b) to train a deep neural network to infer the mean, per-generation dispersal distance from a single population sample of single nucleotide polymorphism (SNP) genotypes, e.g., whole genome data or RADseq data. We show that `disperseNN` is more accurate than two existing methods (Rousset, 1997; Ringbauer et al., 2017), particularly for small to moderate sample sizes, or when identity-by-descent tracts cannot be reliably inferred. After exploring potential shortcomings of our method, we demonstrate its utility on several empirical datasets from a broad range of taxa. The `disperseNN` software is available from https://github.com/kr-colab/disperseNN, where we have also provided a pre-trained model for ease of prediction in new systems.

## Results

### Dispersal estimation using a deep neural network

We use a convolutional neural network (CNN) trained on simulated data to infer parent-offspring distance (Rousset, 1997; Ringbauer et al., 2017) (Figure 1). Concretely, we aim to infer *σ*, defined here as the root­mean-square displacement along a given axis between a randomly chosen child and one of their parents chosen at random (and, we assume directional invariance, so this does not depend on the axis chosen). The CNN takes two pieces of data as input: (1) a genotype matrix, and (2) the distance (e.g. in km) between the two furthest geographic samples. The genotype matrix is put through the network’s convolution layers, while the sampling width is used downstream to convey the physical scale of sampling. The output from the CNN is a single value, an estimate of *σ*. Our software package, `disperseNN`, has several inference-related functionalities: training the CNN on simulated data, predicting *σ* using simulated or empirical data, and pre-processing steps for empirical data. In addition, `disperseNN` includes a pre-trained network that can be used to estimate dispersal without additional training; although the quality of the estimate will depend on how well the data fit the conditions used in training this network; see below for discussion.

**Figure 1:**
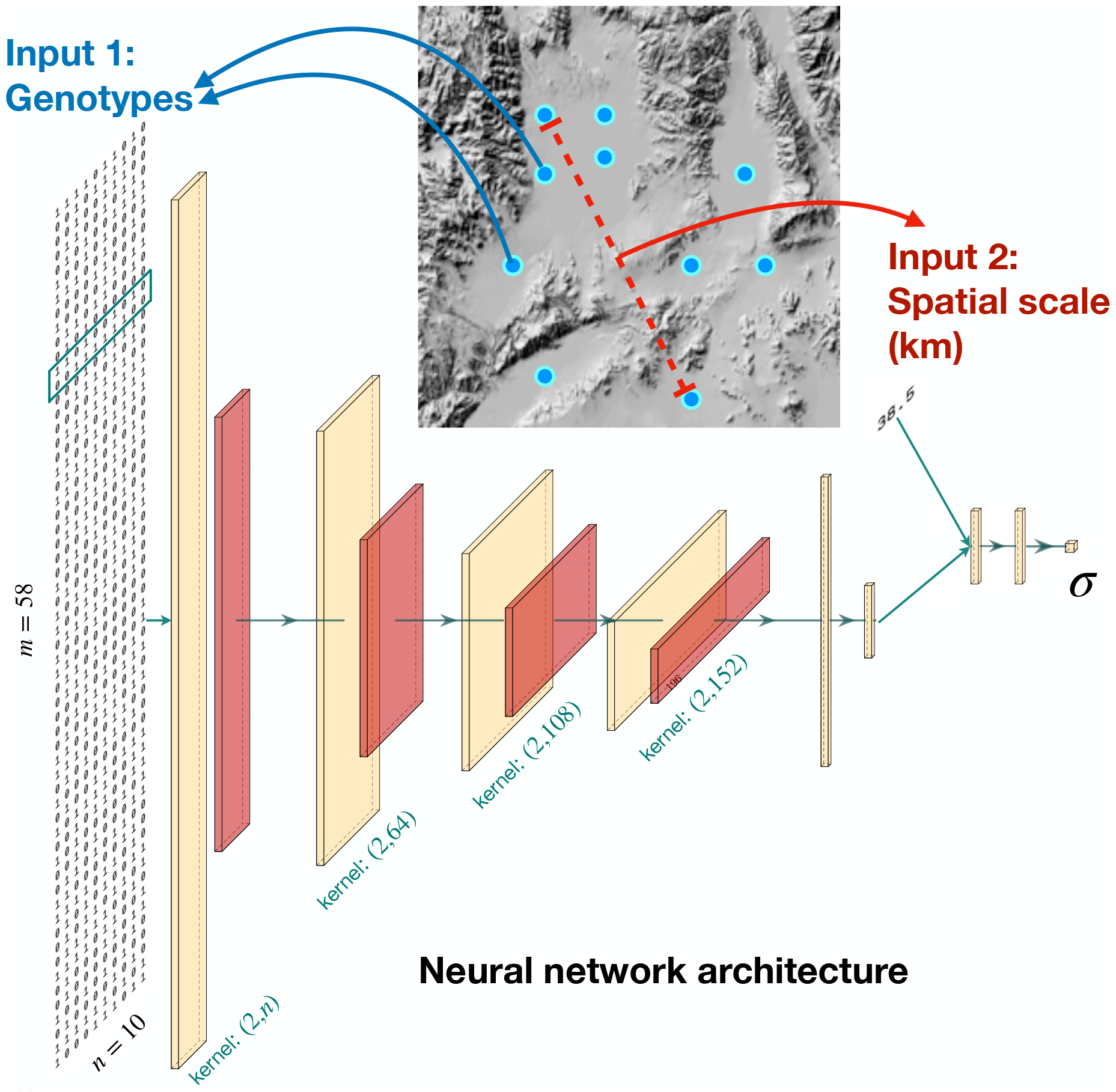
Diagram of the analysis workflow. Blue points are hypothetical sample locations (*n =* 10) on a geographic map. Rectangular tensors are the output from 1D-convolution layers (yellow) and average-pooling layers (red), and the columnar tensors are the output from fully connected layers. The number and dimensions of tensors will vary depending on the input dimensions; this example shows a single haplotype for each individual that is 58 (*m*) SNPs long. The green box over the genotypes shows the size of the convolution kernel for the first layer. The two input branches are eventually concatenated into a single, intermediate tensor. Neural network schematic generated using PlotNeuralNet (https://github.com/HarisIqbal88/PlotNeuralNet).

It is well-known that neighborhood size and dispersal rate not only create genetic patterns of isolation-by-distance, but also influence other facets of population genetic variation. Dispersal affects the site frequency spectrum and other standard summary statistics, such as *θ,* heterozygosity, F*_is_*, Tajima’s D, the variance in *D_xy_,* or nIBS (see, for example, Battey et al., 2020b). Therefore, a natural strategy for estimating *σ* might involve the aforementioned summary statistics, or, as researchers have done for other tasks (e.g., Flagel et al., 2019), use machine learning to extract relevant features from the genotypes themselves in an automated fashion; this is what we seek to do with `disperseNN`.

The convolutional design we use in `disperseNN` is intended to be flexible with respect to input data, as we imagine the use case to be everything from RADseq with minimal, short-range linkage information to whole genome sequence data, where linkage information would be fully preserved. The initial convolutional kernel spans the genotypes of all individuals at once (i.e., a 1-dimensional convolution layer), in a manner that allows the network to glean population-level information at each site, and strides across SNPs two at a time, to make use of correlation patterns when linked SNPs are available. Likewise, we use successive layers of data compression, through convolution and pooling, to coerce `disperseNN` to look at the genotypes at different scales and hopefully to learn the extent of linkage disequilibrium. Counterintuitively, our approach deviates from other dispersal estimators (Rousset, 1997; Ringbauer et al., 2017) because it does not directly utilize the spatial coordinates of sampled individuals, except for calculating the width of the sampling area.

The training data for `disperseNN` are generated using a continuous-space SLiM model adapted from Battey et al. (2020b). Training with `disperseNN` consists of: deciding on training distributions for *σ* and other parameters of the spatial model, simulating training data, and handing the simulation output and targets (true values of *σ*) to `disperseNN` for training the CNN. The analysis pipeline for predicting on simulated data is similar to the training pipeline, while predicting on empirical data involves basic pre­processing of the input data before using `disperseNN` to estimate *σ*. Below, we present findings from several experiments using `disperseNN`, each with its own set of parameters for simulation and training. We describe each experiment briefly in the Results section, and reference different sets of parameters that correspond to each experiment, e.g, “Parameter Set 1”, “Parameter Set 2”, etc. Full details about the different parameter sets are in the Materials and Methods section.

### Comparison with existing methods

We evaluated the accuracy of our method on simulated datasets with a range of *σ* values, using the relative absolute error (RAE) to measure prediction accuracy for each estimate:

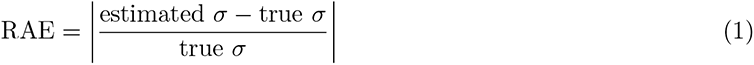

For comparing accuracy between training runs or between methods, we calculate the mean relative absolute error (MRAE) averaged across all test datasets. We found `disperseNN` estimates dispersal rate more accurately than previous genetics-based methods (Figure 2; Parameter Set 1). At small sample sizes (*n* = 10), `disperseNN` was dramatically more accurate than the Rousset (1997) method, the program from Ringbauer et al. (2017) called `IBD-Analysis` run with true identity-by-descent blocks (available from recorded genealo­gies), and `IBD-Analysis` used with empirically estimated identity-by-descent blocks (MRAE values of 0.11, 0.33, 21.41, and 13.6, respectively). Furthermore, the methods other than `disperseNN` produced unde­fined output or convergence errors for 16.4%, 4.6%, and 52.5% of test datasets, respectively. For Rousseťs method, this is due to a negative slope in the least squares fit of genetic distance versus geographic distance, which happens more frequently with a small sample size and larger *σ*. In addition, Rousseťs method and `IBD-Analysis` occasionally produced extreme overestimates for larger *σ* values.

**Figure 2:**
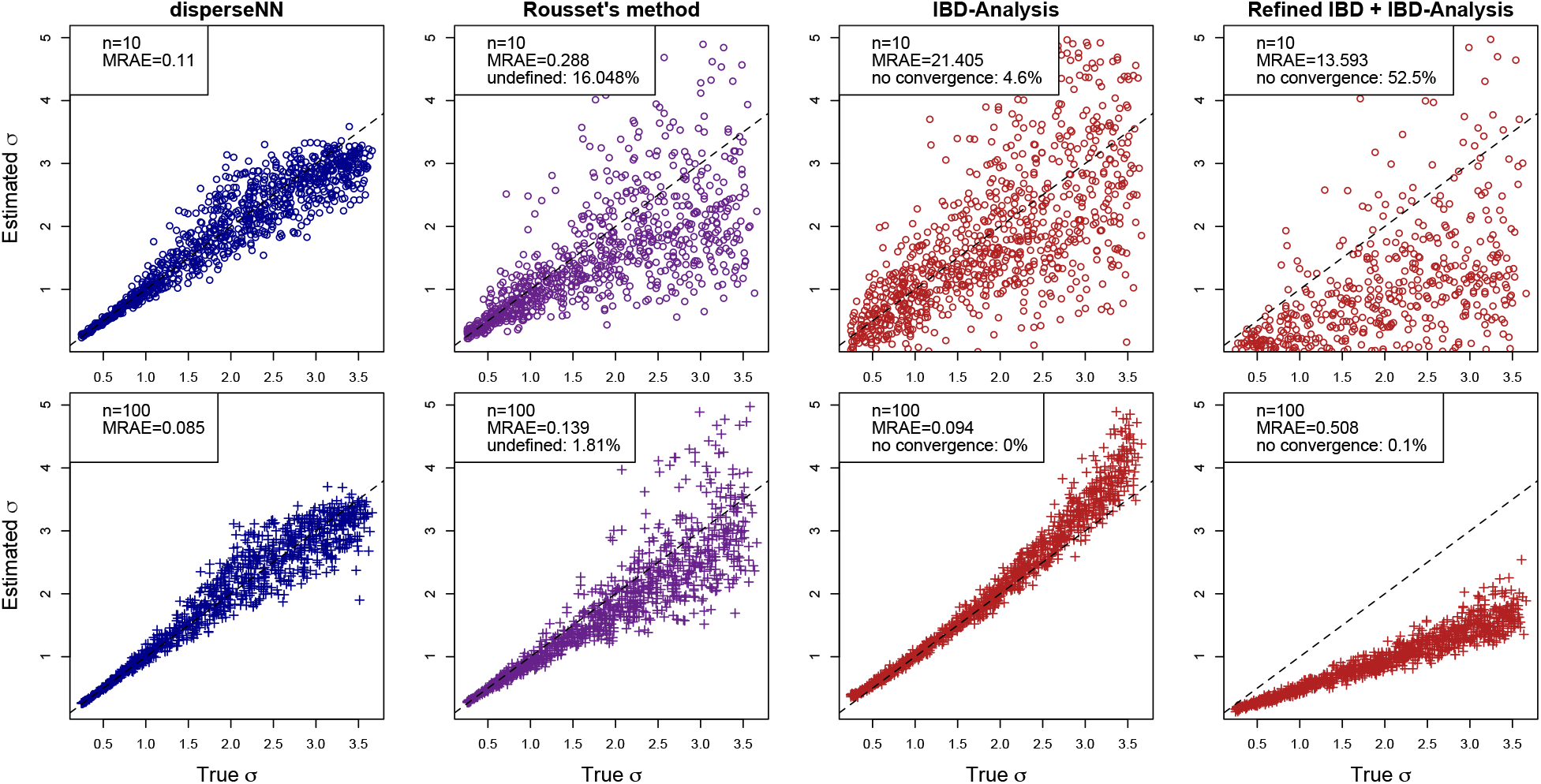
Comparison with existing methods (Parameter Set 1). Here, `disperseNN` is compared with Rousseťs method and the method of Ringbauer et al. (2017) with both true identity by descent tracts (“`IBD-Analysis`”) and tracts inferred by `Refined IBD` (“`Refined IBD` + `IBD-Analysis`”). Two different numbers of sampled individuals are shown: *n* = 10 (top row) and *n* = 100 (bottom row). The numbers of SNPs used with each sample size were 2.5 x 10^5^ and 5 x 10^5^, respectively. The dashed lines are *y = x.* Estimates greater than 5 are excluded from plots but are included in the mean relative absolute error (MRAE) calculation. Methods other than `disperseNN` sometimes produced undefined output; these data do not contribute to the MRAE. The MRAE for `IBD-Analysis` is greater than `Refined IBD` + `IBD-Analysis` with *n* = 10 due to outlier points that inflated the MRAE using the former method and caused undefined output with the latter method.

With a larger sample size (*n* = 100), the accuracy of each method improved to varying degrees, with `disperseNN` and `IBD-Analysis` with true identity-by-descent information performing similarly (MRAE = 0.085, 0.23, and 0.094, respectively). As can be seen in Figure 2, `IBD-Analysis` is extremely accurate with large sample sizes and true identity-by-descent tract information, but had substantially worse performance when tracts must be inferred from genotype data (as would happen in practice). However, despite being underestimated by a constant factor when using empirically inferred tracts, `IBD-Analysis` still seemed to capture a signal of dispersal rate. `IBD-Analysis` with true tracts overestimated *σ* towards the larger end of examined range (`IBD-Analysis`’s bias may be due to limits related to inferring large *σ* relative to the habitat size, or alternatively due to sampling uniformly at random instead of regularly spaced in a grid as in (Ringbauer et al., 2017).) In this comparison `disperseNN` has the advantage of being trained using the true distribution of *σ*; it expects a certain range of *σ* values, which helps it avoid extreme outliers during prediction.

Larger numbers of SNPs improved the performance of `disperseNN` relative to the other methods, although with diminishing returns. See Figures S1 and S2 for results with fewer SNPs. As the number of SNPs decreased, the Rousset method retained good accuracy, better than `disperseNN` for small *σ* values with *n* = 100 samples (although not *n* = 10). For every input size, larger values of *σ* were inferred with correspondingly larger errors (Figure 2), however *relative* error was nearly constant across the range of true *σ* for `disperseNN` (Figure S3). In addition, `disperseNN` slightly underestimated *σ*, and Rousseťs method had larger relative error, when the true value approached the maximum of the examined range. This likely occurs because *σ* can only be so large before approximate random mating is reached; it is difficult to distinguish a large *σ* from a larger-*σ* if both populations have approximately random mating.

### The effect of model misspecification, and how to fix it

A common concern with supervised machine learning methods is that data used for prediction may fall outside of the training distribution. If the training set was simulated with, for example, a small population density, should we expect the trained network to accurately estimate *σ* if the test data come from a species with a large population density? We set out to explore limitations of `disperseNN` using deliberately misspecified simulations, including out-of-sample (i) population density, (ii) ancestral population size, (iii) habitat size, and (iv) restricted sampling area relative to the full habitat. We individually address each scenario by augmenting the training set, which ultimately allows us to produce a trained network that performs well across a wide range of these parameter values. This is important because in practice we often do not have precise estimates of these parameters.

First, we obtained a baseline level of accuracy by training `disperseNN` on data where all simulation parameters were fixed except for *σ* (Parameter Set 2). This resulted in an MRAE of 0.12, averaged across test data whose values of *σ* were drawn from the same distribution as the training set (and other parameters were the same). We next used the model trained on Parameter Set 2 to estimate *σ* in test data where one of the aforementioned variables is drawn randomly from a distribution, and thus misspecified (i.e., differing from the value used for training simulations) to varying degrees (Parameter Sets 3, 5, 7, 9). Such model misspecification reduced the accuracy of *σ* estimation (Figure 3, first column of plots). This reduction in accuracy was most pronounced for misspecified population density and habitat width (MRAE = 0.36 for each). Note that effective population density is an emergent property of the simulation that depends on many aspects of demography; in this experiment we only varied “carrying capacity” (a parameter that affects survivorship), but we expect demographies that are misspecified in other ways (e.g., varying fecundity) to show qualitatively similar results. As shown in Figure 3, the other scenarios also increased error, although more moderately. When a fixed habitat width was assumed, 23% of predictions were larger than the maximum *σ* from training; for other nuisance parameters all predictions fell within the range of *σ* used in training. Rousseťs method suffered from similar increases in error when run on the same misspecified test data as `disperseNN` (Table 1). `IBD-Analysis` was robust to demographic history, consistent with the findings of Ringbauer et al. (2017), however it had issues with the smaller population densities encountered in Parameter Set 3.

**Table 1:** Mean relative absolute errors (MRAE) from misspecification experiments using three different inference methods. `disperseNN` was trained using Parameter Set 2 and applied to test data from the parameter set listed in the second column. The Rousset and `IBD-Analysis` methods were run on the same test data as `disperseNN`, except `IBD-Analysis` used true identity-by-descent blocks; thus, the MRAE is not comparable between `IBD-Analysis` and the other methods which used only 5,000 SNPs. Two values are shown for `disperseNN`: error with misspecified (left) or correctly specified (right) model; the latter was trained while varying the corresponding nuisance parameter, or trained with a sampling strategy and habitat shape that reflect the test data. (*This experiment used a smaller carrying capacity than the others, therefore the `disperseNN` “baseline” MRAE for variable habitat size was 0.09 instead of 0.12. **This experiment used a smaller sample size, *n* = 91, which led to a baseline MRAE of 0.13.)

| Treatment | Parameter Set | **disperseNN** | Rousset | IBD-Analysis |
| --- | --- | --- | --- | --- |
| Baseline | 2 | 0.12 | 0.14 | 0.09 |
| Density | 3 | 0.36, 0.12 | 0.55 | 0.20 |
| Demographic history | 5 | 0.29, 0.12 | 0.39 | 0.10 |
| Habitat size | 7 | 0.36, 0.09* | 0.44 | 0.11 |
| Sampling area | 9 | 0.32, 0.14 | 0.46 | 0.16 |
| Point sampling | 2 | 0.20, 0.12** | 0.59 | 0.29 |
| Asymmetric sampling | 2 | 0.14, 0.12 | 0.19 | 0.10 |
| Transect sampling | 2 | 0.25, 0.09 | 0.20 | 0.09 |
| Hourglass habitat | 2 | 0.33, 0.09 | 0.69 | 0.17 |
| C-shaped habitat | 2 | 0.22, 0.09 | 0.51 | 0.20 |

**Figure 3:**
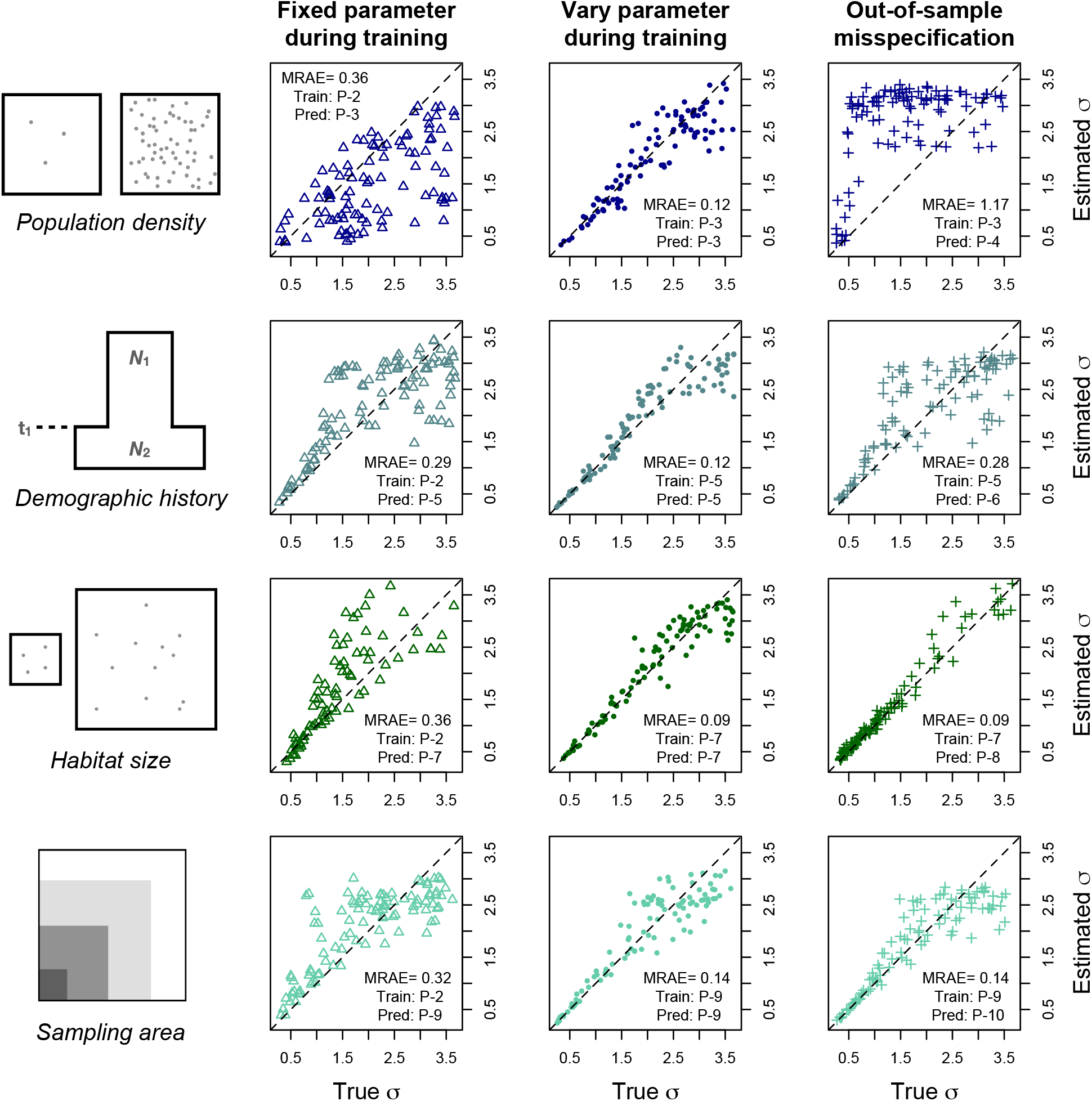
Column 1. Cartoons of unknown parameters that may lead to model misspecification. Column 2. The unknown parameter was fixed during training, but testing was performed on data with different values of the parameter. Column 3. The unknown parameter was *varied* during training, and testing was performed on data from the same distribution. Column 4. The unknown parameter was varied during training, but testing was performed on out-of-sample values, i.e., larger values than were seen during training. The dashed lines are *y = x.* Outliers greater than 3 are excluded from the fixed-habitat-size plot. “Train: P” and “Pred: P” refer to the Parameter Sets used for training and testing, respectively. MRAE is the mean relative absolute error. All analyses used samples of *n* = 100 individuals. (*The third row has a separate baseline MRAE, 0.09, due to using a smaller carrying capacity, which was chosen to alleviate computation time.)

Having observed the effect of misspecification, we next assessed how well the problem could be ameliorated by training across a range of plausible values instead of at a single, fixed value. To do this, for each of the four parameters we train a model using simulations in which the parameter is drawn from a distribution (and, in fact we reuse Parameter Sets 3, 5, 7, 9 for this purpose). In each case, `disperseNN` learned to accurately estimate *σ* when individual nuisance parameters were unknown, with error levels approaching the original MRAE (Figure 3, column 3; Parameter Sets 3, 5, 7, 9). To reiterate, in this procedure we carried out four experiments, varying a single unknown parameter at a time, not in combination. Essentially by treating each unknown parameter as a nuisance parameter during training, the model can become agnostic to the unknown parameter—or else learn a representation for the parameter such that *σ* can be calculated conditional on the learned parameter. This ability is critical for applying supervised learning methods for estimating *σ* where model parameters other than *σ* are unknown.

Although `disperseNN` was able to predict *σ* after including variation in each nuisance parameter in the training set, we next show that extrapolation is limited in some cases for unfamiliar parameter values, i.e., values outside of the distribution used for training. In the preceding trial the same distribution was used for both training and prediction. Next we assessed `disperseNN`’s ability to extrapolate at very large values of each nuisance parameter (Parameter Sets 4, 6, 8, 10), beyond the range used in training (Parameter Sets 3, 5, 7, 9). Results from this experiment were varied (Figure 3, rightmost column): predictions at out-of­sample values of density and ancestral population size were unreliable, but we were able to predict at large, out-of-sample habitat sizes and sampling areas quite well. It is noteworthy that using very large habitat sizes resulted in only a single estimate being 1% larger than the maximum *σ* from training.

Another variable that affects the performance of `disperseNN` is sampling strategy (Table 1; Figure S4). While in the preceding experiments, genotypes were obtained from individuals sampled uniformly at random, real data rarely approximate a uniform sample. We found that after training with uniform sampling, `disperseNN` predictions were somewhat less accurate when given data sampled in clusters or sampled more heavily from one half of the habitat. More substantial error was introduced with transect sampling. How­ever, once `disperseNN` was trained using each of the alternative sampling strategies, accuracy was restored to baseline levels or became even more accurate than uniform sampling (Table 1). In light of this, we advise practitioners to build the empirical sampling strategy into the training simulations for `disperseNN`. In addi­tion, if the shape of the habitat is misspecified during training, predictions from `disperseNN` and the other methods will be skewed (Figure S5; Table 1); however, note that this scenario has additional complexity, because it also includes misspecified habitat *area* (i.e., population size) and misspecified sampling distribution. It is important to consider these complications when designing training simulations for `disperseNN` to ensure accurate predictions. However, unlike existing methods, `disperseNN` has the potential to address these issues during training.

### Training with several unknown parameters

We next sought to train a network that could estimate *σ* even when multiple nuisance parameters are unknown. The resulting network is what we refer to as “the pre-trained network”. To do this, we used large ranges for parameters that control: (i) dispersal distance, (ii) population density, (iii) ancestral population size, (iv) timing of population size change, (v) habitat size, and (vi) the size of the sampling area relative to the full habitat (Parameter Set 11). Furthermore, we exposed `disperseNN` to a range of different sample sizes between 10 and 100 by padding the genotype matrix out to 100 columns during training. However, in all cases sampled individuals were chosen uniformly at random. Training simulations used 5,000 SNPs sampled from a single 100 megabase chromosome with recombination rate 10^-8^: this approach resembles a RADseq experiment, as the loci are spaced out on the chromosome and may be considered mostly unlinked. Last, we collapsed the diploid genotypes output by SLiM into unphased genotypes: 0s, 1s, and 2s; representing the count of the minor allele at each variable site. Through validation with held-out, simulated data, we found that the final model could predict *σ* reasonably well across a wide range of nuisance parameter values (MRAE=0.55; Figure 4). For comparison, we ran Rousseťs method (with density misspecified as *D* = 1, which is close to the mean of the training simulations) and `IBD-Analysis` (with true identity by descent blocks) on the test data from Parameter Set 11, which gave MRAE=32.26 with 24% undefined and MRAE=2.44 with 2% undefined, respectively.

**Figure 4:**
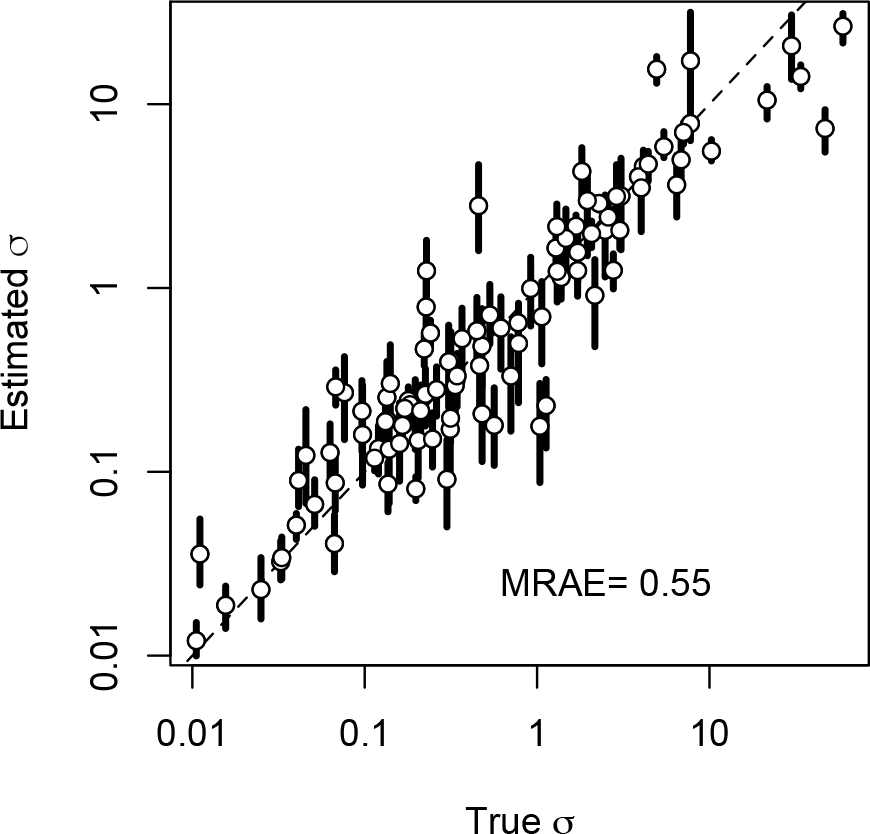
Validation of the pre-trained model (Parameter Set 11). Shown are 100 test datasets, each generated from an independent simulation. Open points indicate the mean estimate from 1000 subsamples of 5,000 SNPs drawn from each dataset, with the sample size varying uniformly between 10 and 100 for each subsample. Also depicted is the range of estimates from the middle 95% of subsamples. The dashed line is *y = x.* Note the log scale. MRAE is the mean relative absolute error.

We provide the learned weights and biases from the above pre-trained network for download as part of the `disperseNN` package. The pre-trained network can be used to quickly estimate *σ* from various species or simulated datasets without additional training or simulations. We note that the pre-trained network for `disperseNN` could in addition be an excellent starting place for transfer learning (Weiss et al., 2016) for specific organisms, sampling designs, or perhaps alternative datatypes (e.g., microsatellite mutations). On our system, the pre-trained network took 6.5 seconds to estimate *σ* using a dataset of 10 individuals and 5,000 variants, with the majority of computation time spent loading software libraries and pre-processing the genotype matrix.

The pre-trained model will be more appropriate for some datasets than others. First, the model was trained on 10 to 100 individuals sampled across a region of known width. Therefore, data collected from a single location are not expected to give accurate predictions (unless the breeding locations were known and spatially distributed). While `disperseNN` can be trained with any number of SNPs, the pre-trained network uses 5,000. Therefore, if fewer than 5,000 variants are available, as in some RADseq datasets, then a new network must be trained to match the empirical number. Padding the input genotypes with zeros will not suffice for using fewer SNPs, as we did not train with zero-padding. Although we aimed to produce a pre-trained model that is widely applicable, many of the attempted simulations either resulted in population extinction, or could not be simulated due to computational constraints, which resulted in parts of parameter space not represented in the realized, multivariate training distribution (Figure S6). Therefore, we expect this model to be most applicable for smaller populations that fall solidly inside of the training distribution. See Parameter Set 11 in the Materials and Methods section for “prior” ranges.

Additional training will be beneficial in some situations. If independent estimates for nuisance parameters or better-informed “prior” ranges are available, new training data may be tailored using the better-informed values. Species range maps with detailed geographic boundaries can be simulated with SLiM (since version 3.5) or `slendr` (Petr et al., 2022), which could be superior to the square map we used. Importantly, if empirical parameters fall outside of the training distributions used for the pre-trained network, e.g., very large sampling area, then new training data will need to be generated that reflect the real data.

### Quantifying uncertainty

In addition to helping us generate training data, simulation also allows us to quantify uncertainty through validating our models on held-out test datasets. Indeed, our reported values of MRAE give a sense of how much error to expect when applying the method to real data, in so far as the data resemble a typical draw from our test simulations. For example, in the above experiments that included one or zero nuisance parameters, the MRAE from in-sample tests was on the order of 0.12. Therefore, using a model with MRAE of 0.12 we might expect future predictions to be off from the true values by about 12%. However, if the real data are not well represented by the simulations, for example if the density of the analyzed population does not resemble that of the training simulations, then predictions might be less accurate, or biased.

Since we get distinct estimates for each subset of *m* SNPS, we can also assess uncertainty by looking at the range of variation among these estimates, i.e., through non-parametric bootstrapping. Each subsample of *m* SNPs from the same set of sampled individuals gives a different estimate of *σ* because of the varying genealogical histories that underlie different subsets of genomic loci, so the range of variation reflects the uncertainty arising from this genealogical noise. However, note that the bootstrapped estimates are not independent, because of linkage and because they come from a single set of individuals. The `disperseNN` program provides a built-in functionality for performing this bootstrapping procedure, and will report the distribution of estimates across replicate draws of *m* SNPs (each draw is made without replacement from the complete set of available SNPs, but the replicates are drawn independently and so may overlap).

Although the distribution of these estimates should reflect uncertainty somehow, it is not immediately clear how to convert this into a formal quantification of uncertainty. This distribution of estimates is not a sample from a well-calibrated posterior distribution (nor should we expect it to be): in the test data for the pre-trained model (Figure 4), the true *σ* was covered by the middle 95% range from the bootstrap distribution for only 51% of simulated datasets. However, we can inflate the interval obtained by a scalar value such that our bootstrap interval is better calibrated. On our validation set for the pre-trained model this scalar value is 3.8, which leads to intervals that cover the true value for 95% of our test simulations. (If 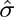 is the mean of the bootstrap estimates, and *a* and *b* are the 2.5% and 97.5% quantiles, respectively, then the resulting interval is from 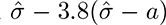 to 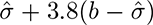.) However, if this is to be a recipe for a well-calibrated credible (or, confidence) interval, then it needs to apply regardless of the situation: i.e., the magnitude of the error should be a roughly constant multiple of the range of the bootstrap estimates. Happily, this seems to be the case: in our experiments, we found the error to be roughly a constant multiple of the width of the range of bootstrap estimates. Concretely, if *σ* is the true value, 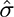 is the estimated value, and *w* is the range of values from 100 bootstrap estimates, then 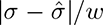 has no significant associations with any of the model parameters using Parameter Set 11; see Figure S7.

In summary, this suggests that the middle 95% interval of bootstrap estimates, inflated by a factor of 3.8, can stand in for a 95% credible interval for results obtained from our pre-trained neural network. Of course, since this is an empirically derived result, we cannot guarantee the same inflation value to be appropriate for other networks or for datasets not well-represented by the simulations in the training set for our pre-trained model.

### Empirical findings

We used `disperseNN` to estimate *σ* from a diverse set of organisms using preexisting empirical datasets. The pre-trained `disperseNN` model works with a wide range of genetic data, including low coverage whole genome sequencing or RADseq data, because genotypes were not phased during training. Deviations in mutation rate between the training and empirical data will not affect the results, because we trained `disperseNN` with a fixed number of SNPs—5,000—sampled throughout the genome. Likewise, deviations in recombination rate are unlikely to be a concern because we created the pre-trained model with SNPs that are fairly spaced out across the genome and are mostly independent, and the empirical SNPs are similarly distributed. For some empirical datasets, we analyzed a subset of sample localities in order to keep the sampling width less than 1,000 km; accordingly, we report sample sizes and sampling widths from the subsampled region, rather than the full dataset. Also, we ensure that each sample location is represented by only one individual. For each dataset, we used `disperseNN` to prepare the two inputs for the CNN: the SNP table was converted to a genotype matrix and the distance between the furthest individuals was calculated. Next, we used `disperseNN` to predict *σ* on each of 1,000 independent bootstrapped samples of 5,000 SNPs, obtaining a distribution of *σ* estimates. Table 2 shows the mean and approximate 95% credible interval of *σ* estimates, along with other analysis parameters, for each empirical dataset. We note, however, that the credible intervals for empirical datasets with smaller numbers of SNPs may be poorly calibrated (e.g., *Peromyscus leucopus,* that has only slightly more than 5,000 SNPs).

**Table 2:** Empirical results. The σ column is the mean from 1000 subsamples of 5,000 SNPs. “95% CI” is the credible interval obtained from bootstrapping. The “Previous” column shows previously published estimates for dispersal distance. N_loc_ is the neighborhood size using the Rousset calculation. In other columns, n is sample size, S is the width of the sampling area in kilometers, and “M. dist.” is the Mahalonobis distance from the center of the training distribution with respect to five summary statistics: nucleotide diversity, Tajima’s D, inbreeding coefficient, observed heterozygosity, and expected heterozygosity.

| Species | Common name | Region | $\sigma$<br>(km) | 95% CI<br>(km) | Previous<br>(km) | $N_{loc}$ | $n$ | $S$<br>(km) | M. dist. |
| --- | --- | --- | --- | --- | --- | --- | --- | --- | --- |
| *Zosterops borbonicus* | Réunion grey white-eye | Réunion | 4.97 | (1.76, 13.83) | NA | 295 | 41 | 62 | 4.59 |
| *Peromyscus leucopus* | white-footed mouse | New York | 0.77 | (0.32, 1.67) | 0.03-0.11 | undef. | 12 | 38 | 8.15 |
| *Anopheles gambiae* | African malaria mosquito | Cameroon | 10.29 | (2.00, 48.03) | 0.04-1.57 | 52 | 29 | 278 | 9.62 |
| *Bombus bifarius* | two-form bumble bee | Washington | 14.75 | (5.60, 37.28) | 1.2-5 | 1,147 | 14 | 273 | 10.47 |
| *Bombus vosnesenskii* | yellow-faced bumble bee | California | 7.70 | (1.21, 38.11) | 1.2-5 | 3,944 | 18 | 169 | 11.83 |
| *Hippoglossus hippoglossus* | Atlantic halibut | Canada | 4.29 | (0.71, 33.85) | NA | undef. | 11 | 193 | 14.59 |
| *Crassostrea virginica* | eastern oyster | Canada | 1.52 | (0.72, 4.31) | 21.9 | 1,435 | 13 | 187 | 19.69 |
| *Canis lupus* | grey wolf | N. America | 15.68 | (2.36, 107.3) | 98-147 | 35 | 13 | 721 | 25.42 |
| *Helianthus petiolaris* | prairie sunflower | Kansas | 1.00 | (0.39, 3.52) | 0.156 | 9 | 11 | 204 | 45.28 |
| *Zosterops olivaceus* | Réunion olive white-eye | Réunion | 1.05 | (0.27, 4.36) | NA | 2,392 | 10 | 50 | 45.97 |
| *Helianthus argophyllus* | silverleaf sunflower | Texas | 1.04 | (0.38, 4.08) | 0.156 | 57 | 30 | 307 | 86.49 |
| *Arabidopsis thaliana* | thale cress | Spain | 1.36 | (0.28, 5.05) | 0.001 | 35 | 35 | 80 | 198.25 |
| *Arabidopsis thaliana* | thale cress | Sweden | 0.44 | (0.20, 0.93) | 0.001 | 84 | 84 | 325 | 428.17 |

When available, we report previous dispersal estimates from the literature. Independent estimates came from a variety of methods including mark-recapture, tracking devices, and the Rousset method. Overall we find a correlation (*r*^2^ = 0.39; *p =* 0.03; linear regression on log-transformed inputs) between our estimates and previous estimates using different methods. We might expect each of the analyzed empirical datasets to deviate from our training set in some way. To get a rough estimate of the “distance” between an empirical dataset and our training set we calculated five summary statistics—nucleotide diversity, Tajima’s D, *F_IS_* (an estimate of inbreeding), observed heterozygosity, and expected heterozygosity—and calculated the Mahalanobis distance between the centroid of the training distribution and each dataset, according to: *D^2^ =* (*x – m*)*^T^*· *C*^-1^ · (*x – m*), where *D^2^* is the Mahalanobis distance squared, re is a vector of summary statistics from an empirical dataset, and *m* and *C* are the vector of means and the empirical covariance matrix of the summary statistics in the training data. Thus, smaller distances have summary statistics more similar to the training distribution, and distances larger than roughly 40 fall outside of the training distribution (Figure S8). However, a small Mahalanobis distance does not guarantee that the model is well-specified for a given dataset. In fact, we expect error to accumulate due to misspecified sampling scheme, irregular habitat, and other variables that affect population genetic variation, some of which are mentioned below.

#### Zosterops

Réunion grey white-eye and Réunion olive white-eye are endemic to the island of Réunion with approximate land area of 2500 km^2^. These species’ restricted ranges make them ideal for analyzing with our pre-trained model. We analyzed the RADseq data from Gabrielli et al. (2020) including 41 individuals and 7,657 SNPs from *Z. borbonicus* and 10 individuals and 6,103 SNPs from *Z. oliυaceus.* Our estimate for *Z. borbonicus* was 5.0 km, however the estimate in *Z. oliυaceus* was smaller, 1.1 km. Although we are not aware of other dispersal estimates in these species, the data curated by Paradis et al. (1998) include natal dispersal estimates for 75 birds, and the smaller species, comparable in size to *Zosterops,* have dispersal distances in the range of 1-20 km. The mean estimates for both species fall within the range from Paradis et al. While the data for both *Zosterops* species are similar, summary statistics in *Z. oliυaceus* were further from the centroid of the training distribution.

#### Peromyscus leucopus

From the white footed mouse RADseq dataset of Munshi-South et al. (2016) we analyzed 12 individuals collected from the New York City metropolitan area, with 5,536 SNPs. We estimated dispersal distance to be 770 m. For comparison, Keane (1990) and Jacquot and Vessey (1995) measured natal dispersal in white footed mice in rural locations. They reported mean dispersal of 85-109 m in males and 25-88 m in females, which are smaller than our estimate. However, their estimates are likely constrained to some degree by the small study areas used for recapture. Indeed not all mice were recaptured in Jacquot and Vessey (1995), leaving open the possibility of long distance movements outside of the study area. For example, Murie and Murie (1931) documented travel distances greater than 1 km in *Peromyscus maniculatus.* Occasional long distance dispersal may help reconcile the difference between previous estimates and ours.

#### Anopheles gambiae

From the whole genome resequencing dataset from the *Anopheles gambiae* 1000 Genome Consortium (2021) we analyzed 29 individuals with 11 million SNPs. Our estimate in *A. gambiae* of 10.3 km is substantially larger than mark-recapture estimates. For comparison, Epopa et al. (2017) measured individual *A. coluzzii* dispersal distances between 40 to 549 m over seven days; however the geographic study region was restricted to a single village. Another study, Gillies (1961), reported mean dispersions of 837 to 1,577 m, depending on sex and release location. A third study reported *per-day* movements of 350-650m (Costantini et al., 1996). It is unclear to what degree long-distance dispersal in mosquitos contributes to effective dispersal and gene flow. Remarkably, the recent study of Huestis et al. (2019) captured *A. gambiae* and other mosquito species 40 m to 290 m above the ground, suggesting a wind-borne dispersal mechanism. Assuming average wind speeds, Huestes et al. estimated that each year tens of thousands of *A. gambiae* individuals migrate 10s or 100s of km in the atmosphere of the studied region. These findings suggest that dispersal potential in this species is considerably larger than once thought. Significant long-range dispersal in *A. gambiae* is consistent with some predictions in the species, as there is little genetic differentiation across portions of the species range (e.g., West Africa), while at broader scales structure is appreciable (Anopheles gambiae 1000 Genome Consortium, 2017).

#### Bombus

From the dataset of Jackson et al. (2018) we examined RADseq data from two bumble bee species, *B. bifarius* and *B. vosnesenskii* with samples sizes of 14 and 18, and 8,073 and 6,725 SNPs, respectively. Our estimated dispersal distances were 14.8 km and 7.7 km for the two. These species are eusocial, so our dispersal estimate should reflect the distance traveled by queens that start successful nests. Mark-recapture analyses have found a minimum distance traveled by queens in other *Bombus* species of 1.2 km (Carvell et al., 2017), and using genetic full-sib reconstruction resulted in 3-5 km (Lepais et al., 2010). These estimates are particularly relevant, as they measure natal dispersal from the birth location of the queen. Even so, these values represent a lower bound distance that queens disperse, as there was potential for longer-distance dispersal events that fall outside of the study area. Our results may offer a glimpse into bumble bee dispersal including longer distances that would be difficult to measure directly.

#### Hippoglossus hippoglossus

From the RADseq data of Kess et al. (2021) we analyzed 11 individuals with 69,000 SNPs. Tagging studies find mean halibut movements greater than 100 km (Liu et al., 2019). However, the distance traveled by adults in search of food may be considerably larger than the quantity we wish to estimate which is proportional to the mean distance between birth location and parental birth location. Indeed, there is spatial structure distinguishing Atlantic halibut stocks due to spawning site fidelity (Shackell et al., 2021). Therefore, the observed sample locations—used to calculate the second input to `disperseNN`— are likely foraging locations that may differ significantly from the breeding locations. However, if assumptions about the size of the spawning area can be made, `disperseNN` provides a novel approach for inferring effective *σ* in foraging or migrating individuals for whom “home” locations are not known. Our estimate of 4.3 km (using the sampling width as the second input) could be close to the true dispersal distance if birth site fidelity is quite high. In another large marine species, *Diplodus sargus sargus,* natal dispersal distance was measured to be 11 km using otolith chemistry (Di Franco et al., 2012).

#### Crassostrea virginica

From the RADseq data of Bernatchez et al. (2019) we analyzed 13 individual eastern oysters with 7,097 SNPs. This species has larval dispersal (Vercaemer et al., 2010) and occasional adult translocations (Bernatchez et al., 2019). Our estimate of 1.5 km is much smaller than the previous estimate of 21.9 km (Rose et al., 2006). We offer several possible explanations for this discrepancy. We expect that oyster dispersal is at least in part passive, particularly during the larval stage, and so depends on local currents. The previous estimate was from a different sample region, Chesapeake Bay, which likely has different local conditions than the coast of Canada where the samples that we analyzed were collected. Second, the previous estimate used microsatellite loci to estimate density in order to implement the Rousset method. Density is notoriously difficult to estimate from genetic data, so it would not be surprising if this step contributed to error. In contrast, `disperseNN` is designed to work around the unknown density parameter. However, we note that the range of this species is likely to be closer to a narrow strip than the broad square or rectangle used in our simulations.

#### Canis lupus

From the RADseq dataset of Schweizer et al. (2016) we analyzed data from 13 individual wolves genotyped at 22,000 SNPs. Exceptionally good data exist on wolf dispersal from radio collars. A commonly reported value for this species is the distance traveled by adults that disperse between territories. For example, some estimates for this value include 98.1 km (Jimenez et al., 2017), 98.5 km (Kojola et al., 2006), and 147.0 km (Barry et al., 2020). However, not all individuals disperse from their natal territory. For example, Kojola et al. (2006) and Barry et al. (2020) reported that 50% and 47% of individuals dispersed between territories, respectively. Jimenez et al. (2017) reported more nuanced statistics: 18% of collared individuals had documented dispersal, survival was lower in dispersers, and not all dispersers reproduced. It is unclear how frequent breeding occurs *within* the natal pack; if 85-90% of reproduction occurred without movement between territories, then our estimate of 15.7 km might be reasonably close to the true, effective dispersal distance.

#### Helianthus

We analyzed two wild sunflower species from Todesco et al. (2020): *Helianthus petiolarus* (*n =* 11; 61,000 SNPs) and *H. argophyllus* (*n* = 30; 60,000 SNPs), with whole genome resequencing data. Wild sunflowers regularly outcross, therefore the estimated *σ* in part reflects pollinator distance, in addition to transport of seeds by animals and other methods. Previously, Arias and Rieseberg (1994) reported the frequency of hybridization between cultivated and wild sunflowers at distances between 3 m and 1000 m; if we convert these hybridization-frequencies to counts of hybridization events, the mean distance of these pollination events was 156 m. The estimates from `disperseNN` were larger: 1000 m and 1040 m in *H. petiolaris* and *H. argophyllus,* respectively. These estimates seem large but may be reasonable if pollination occurs via bees, which can have foraging ranges greater than 1 km (Osborne et al., 2008; Visscher and Seeley, 1982). Studying foraging distance in pollinators is an active area of research, however Pasquet et al. (2008) used an exceptionally large study area and radio trackers to find a median flight distance of 720m in carpenter bees. Our estimates for the two analyzed *Helianthus* species were similar to each other.

#### Arabidopsis thaliana

From the whole genome resequencing dataset of the The 1001 Genomes Consortium (2016) we analyzed two sampling clusters from different geographic regions: Spain (142,000 SNPs, n=35) and Sweden (124,000 SNPs, n=84). Our *σ* estimates from these two groups of samples were 1,360 m and 440 m, which are considerably larger than the average distance that seeds fall from the parent plant; Wender et al. (2005) estimated that the average distance traveled by *A. thaliana* seeds with wind is less than 2 m. However, occasional long distance seed dispersal, e.g., via water or animals, and infrequent outcrossing via insect pollination may inflate the effective dispersal distance in this species. The outcrossing rate of *A. thaliana* has been estimated to be 3 x 10 ^3^ (Abbott and Gomes, 1989). Importantly, *A. thaliana* is predominantly selfing and the analyzed samples are (naturally) inbred, while our training set which did not include selfing. *A. thaliana* has experienced a known population expansion (Tyagi et al., 2016), and although we attempted to account for demographic history during training the true history of *A. thaliana* may not be well-represented by our simplistic range of population histories. There was a three-fold difference in estimated dispersal distance between the analyzed populations, perhaps due to local environmental differences between Spain and Sweden or different pollinator species.

## Discussion

### Dispersal estimation using deep learning

Understanding how organisms move across land or seascapes is critical for gaining a full picture of the forces shaping genetic variation (Wright, 1943; Kimura and Weiss, 1964; Barton et al., 2002). However, it remains difficult to confidently infer spatial population genetic parameters. Here we present a deep learning framework, `disperseNN`, for estimating the mean per-generation dispersal distance from population genetic data. There are several advantages of our method over existing population-genetics-based estimators, including improved accuracy for small (*n =* 10) to moderate (*n =* 100) sample sizes, accessible input data (unphased SNPs), and the ability to infer dispersal distance in the face of unknown model parameters such as population density. It is unclear how competitive `disperseNN` would be with other programs if given very large sample sizes (e.g., thousands of genotypes), because the memory requirements for such a scale may be limiting. These improvements open the door for using DNA to infer dispersal distance in non-model organisms where population density is unknown or identity-by-descent tracts are out of reach. Perhaps more importantly, because `disperseNN` uses a form of simulation-based inference, analyses do not depend on idealized mathematical models of flat, featureless space, and can be tailored for the particular study system, for instance detailed habitat maps and independent estimates for key model parameters can be readily incorporated.

Unlike previous genetics-based estimators that use geographic distances between individuals, our neural network does not see the relative spatial locations of individuals. This means that our neural network could in theory be applied to genetic data for which sampling locations are unavailable due to ethical or legal protections, or applied to adult individuals that have ranged far from their nesting or spawning area. However, to do so an estimate of the sampling width is required as input by `disperseNN`. We had initially sought to convey the spatial sampling coordinates to the network, inspired by Rousset (1997) and Ringbauer et al. (2017), however doing so did not improve the model’s accuracy. This surprising result may have to do with limitations of the chosen architecture. Or, perhaps the spatial coordinates convey little additional information for the task we are interested in beyond the signal inherent to the genotypes. `disperseNN` is able to infer dispersal without individual sample locations, because the dispersal rate affects not only pairwise genetic distances between individuals, but also population genetic variation more generally, such as the site frequency spectrum and its transforms (e.g. nucleotide diversity), inbreeding coefficients, and linkage disequilibrium (Battey et al., 2020b). Additionally, it is well-known that genotypes can often be used to obtain a reasonably good estimate of relative sampling locations (e.g., by PCA; Novembre et al., 2008). `disperseNN`, by using a convolutional neural network with a genotype matrix as its input, is able to capture population genetic information from raw data as has been seen in a few prior contexts (e.g., Flagel et al., 2019; Sanchez et al., 2021; Gower et al., 2021).

Another strength of the deep learning approach is its versatility. In particular, `disperseNN` can be used with unphased SNPs and small sample sizes, which makes it applicable for a variety of genomic dataset types. In contrast, recently developed tools for dispersal estimation require identity-by-descent blocks as input (Ringbauer et al., 2017; Al-Asadi et al., 2019). Although these methods perform well when high quality data are available, phasing and identity-by-descent inference in non-human genomes is a considerable challenge, especially for RADseq. Unphased SNPs, on the other hand, are more widely available.

Next, `disperseNN` can infer dispersal without *a priori* knowledge of other important parameters, such as population density. In contrast, the commonly used Rousset method requires an independent estimate for population density in order to infer dispersal distance. Our supervised learning approach can learn to predict *σ* in the face of unknown density, which is achieved by exposing the network to training datasets with various densities. Through this procedure, `disperseNN` successfully learned to estimate *σ* in test datasets regardless of density, conditioned on true density being within the training distribution. While that is so, we still observed misspecification for large, out-of-distribution densities, which caused the network to overestimate *σ*. We used the same approach to deal with uncertainty of various other parameters. On the other hand, if independent estimates for some parameters or better-informed “priors” are available, then training can be customized to reflect the known parameters. It is worth mentioning that our general inference framework can be easily modified to infer multiple parameters at once, namely population density and dispersal, although we have not explored this yet.

Thus far we have focused on indirect estimation of dispersal distance, without measurements of how far individuals move. For a review of other genetic techniques for estimating dispersal distance, including direct and indirect methods, see Broquet and Petit (2009). Recently, two studies have used close kin mark­recapture approaches for estimating dispersal distance, which were applied to mosquito species (Jasper et al., 2019; Filipović et al., 2020). Close kin mark-recapture uses the genome of a close relative to represent a “recapture”, thereby skipping the need to physically recapture individuals. These promising new methods estimate dispersal distance by modeling the spatial distribution of close kin. In theory, our approach may offer advantages over close kin mark recapture: `disperseNN` aims to estimate *effective* dispersal, has no requirement for close kin to be captured together, and works with small sample sizes (*n* = 10). The ability to capture kin relies on a sample size that is a sufficiently large proportion of the local population size, which is not always feasible.

### Limitations

Although training on simulated data allows great flexibility, the simulation step was also a limitation for the current study. In particular, generating the training data for our pre-trained network involved very long computational run times and large memory requirements: up to 175 gigabytes of RAM and two weeks of run time for the largest parameterizations of individual simulations. Shortcuts were used to reduce simulation time, including running fewer spatially-explicit generations, and sampling multiple times from each simulated population (see Materials and Methods). Of course, if new training data are generated for a population that is comparatively small, then the simulation burden will be smaller.

As with many statistical approaches, `disperseNN` has limited ability to generalize outside of the range of parameter values on which it was trained. Although exposing the model to training datasets with varying parameter values successfully produced estimates robust to variation in those parameters, the resulting models were still unable to provide good estimates for out-of-sample data. For instance, if the test data came from a population with density higher than those the network had seen during training, *σ* was overestimated. Likewise, prediction error increased if the test data had a larger spatial sampling area than the network saw during training. Therefore we expect the pre-trained model from our empirical analysis to be most accurate for smaller spatial samples from smaller populations—parameters that fall inside the training range—while applications to larger populations may be more questionable. In fact, it is generally recommended to restrict the sampling area to a small region when estimating *σ* to avoid issues with environmental heterogeneity and patchy habitats (Broquet and Petit, 2009; Shipham et al., 2013). However, a larger sampling area is clearly required to infer a large *σ*.

Another potential issue with our approach is complex demographic history. As demographic perturbations leave a footprint in contemporary genetic variation, demography may bias estimates of *σ* for a neural network trained with a particular history, e.g., constant size. This issue is by no means unique to our analysis. Leblois et al. (2004) showed that dispersal rate inference using Rousseťs technique was affected by past demographic values rather than recent population density. We attempted to address this in our analysis, by simulating under random two-epoch models. This approach produced accurate estimates for test data with a similar two-epoch history. However it also suggests that different, more complex demography may reduce accuracy, for example a more extreme bottleneck than was simulated in training, fluctuating size, pulse admixture, or perhaps population structure not captured in our simulations (e.g., barriers to dispersal or range expansion). Identity-by-descent based methods may alleviate the effect of ancestral population structure because long identity-by-descent tracts originate from the recent past (Barton et al., 2013). Similar to demographic history, other model misspecifications such as complex habitats and environmental heterogeneity could also be sources of error for estimation using our method.

Likewise, in our model demography is uniform across space. This assumption may be nearly true—or, at least useful—for certain applications, particularly if the sampling area is small. However, in reality we expect dispersal to vary across space due to a variety of reasons: for example, mountain ranges will prohibit dispersal for many species. Alternatively, suitable habitat is often discontinuous, and dispersal between patches may be different than within patches. Likewise, heterogeneous habitat can generate source-sink dynamics. Existing methods do infer heterogeneous dispersal surfaces across space (Petkova et al., 2016; Al-Asadi et al., 2019), but have limitations including (i) estimating relative differences in dispersal as opposed to the magnitude of dispersal, or (ii) requiring identity-by-descent data as input.

When we included multiple nuisance parameters (Figure 4; Parameter Set 11), the MRAE was larger than that of experiments with only one or zero nuisance parameters (Figure 3; e.g., Parameter Set 3). This difference can be partly explained by the larger number of parameters with potential to confound. In addition, the *range* of values explored for *σ*, as well as for nuisance parameters, were orders of magnitude larger than those of the other experiments. Finally, it is important to note that in our simulations, a single parameter was used for typical “mother-child” dispersal distances, “mother-father” mating distances, and the “interaction distance” over which population regulation occurs. (However, the quantity we estimate is mean parent-offspring distance, which incorporates the first two.) If these quantities are very different in a real population, the pre-trained network may be less reliable than we have estimated.

### Interpretation of empirical findings

We estimated *σ* in a diverse set of organisms using publicly available datasets. These included data obtained by both whole genome shotgun sequencing and RADseq—i.e., variations on standard RADseq (Baird et al, 2008) or genotyping-by-sequencing protocols (Elshire et al., 2011). Rather than simulate scenarios that would be appropriate to each species independently, we trained a single `disperseNN` model designed to estimate *σ* without *a priori* knowledge of density, ancestral population size, or species range.

The majority of empirical results from `disperseNN` were sensible, however our estimates for *A*. *thaliana—* particularly in the population located in Spain—are likely overestimates, in part due to the lack of selfing in our training simulations. *A*. *thaliana* also had levels of heterozygosity and inbreeding that were outside the range of values observed in the training set, a feature reflected in the Mahalanobis distances between training and prediction sets. In the future, `disperseNN` might be better tuned to analyze selfing species, but this would require simulating additional training data and subsequent validation steps.

Our approach led to consistently larger dispersal estimates than mark-recapture experiments. Mark­recapture data were available for three of the analyzed taxa—white footed mouse, *Bombus,* and *Anopheles.* However the mark-recapture estimates for *Anopheles* are not ideal, as they represent adult-travel distances instead of parent-offspring distances. In contrast, the measurements from bumble bees (Carvell et al., 2017) and mice (Keane, 1990; Jacquot and Vessey, 1995) are particularly relevant, as they measure the distance traveled by queen bees from the original hive or individual mice between birth location and adult territory. In all three cases our estimate was larger than the mark-recapture calculation, which suggests either an upward bias in the `disperseNN` output or underestimation in the mark-recapture estimates. In each mark-recapture study the geographic recapture area was smaller than the sampling area we provided to `disperseNN`. It is likely that long-distance dispersers, even if less common, are missed during the recapture step, which would bias the inferred dispersal distance downward in direct, mark-recapture studies.

### Population genetics for spatial ecology

We conclude with a call for increased development and applications of population genetics methods for spatial ecology applications. Dispersal is one of the main factors controlling metapopulation dynamics (Leibold et al., 2004), as well as the total population size and whether a population persists (Gadgil, 1971). Therefore, dispersal estimates are critical for choosing appropriate settings in population viability analyses (Akçakaya and Brook, 2008). Likewise, geographic habitat shifts are ongoing for many species, and species’ survival may thus depend on their ability to disperse fast enough to follow rapidly changing local conditions (Wiens, 2016). Thus, obtaining values for dispersal distance are important for species distribution modeling which is used to project future species ranges (Wiens et al., 2009). In the comprehensive review of Driscoll et al. (2014), the authors present a list of 28 applications for which dispersal values were needed in conservation management, and report several independent calls for improved dispersal information and dispersal inference methods (Broquet and Petit, 2009; Hadley and Betts, 2012; Ceballos et al., 2009; Kingsford et al., 2009; Sutherland et al., 2006; Noss et al., 2009; Pullin et al., 2009).

Characterizing dispersal is also important for managing animal populations relevant to human health. For example, in the fight against malaria we must identify migration corridors and source-sink dynamics in mosquito vector species to allocate pesticide treatment and to predict the spread of genetic variants conveying insecticide resistance (Clarkson et al., 2020). Understanding dispersal is particularly important for modeling and implementing gene-drive strategies (Champer et al., 2021; Beaghton and Burt, 2022; North et al., 2013, 2019, 2020; Beaghton et al., 2016, 2017) for controlling the spread of mosquito-borne diseases including malaria.

Direct methods such as radio tracking or genetic identification may provide near-perfect measurements of dispersal within the generation or generations analyzed. However it is often more valuable to know the expected dispersal distance over many generations, conditional on survival and successful reproduction of the dispersing individuals. For example, the day-to-day foraging distance or seasonal migration distances traveled by adults may differ from the effective dispersal distance. Direct methods such as mark-recapture are often expensive and as a result are limited to relatively small geographic areas, which may ignore long distance movement and bias the resulting estimate. Therefore, population genetic tools may complement direct methods for improving our understanding of dispersal.

## Materials and Methods

### Simulations

Training datasets were simulated using an individual-based, continuous-space model based on the SLiM model used in Battey et al. (2020b). The simulation is initialized with hermaphroditic, diploid individuals distributed randomly on a square habitat. The life cycle of an individual consists of stages for dispersal, reproduction, and mortality. Each offspring disperses from the maternal parent’s location by an independent random displacement in each dimension that is Gaussian distributed with mean zero and standard deviation *σ_f_.* The mate of each individual in each time step is selected randomly, with probability proportional to a Gaussian density with mean zero and standard deviation *σ_m_,* up to a maximum of 3*σ_m_* units in space. The probability of survival of an individual depends on the local population density around the individual, allowing the total population size to fluctuate around an equilibrium. Specifically, the local population density around individual *i* is measured using a Gaussian kernel with standard deviation *σ_c_* as 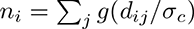, where *g*(*d*) is the standard Gaussian density and *d_ij_* is the distance between individuals *i* and *j* and the sum is only over individuals out to a maximum distance of 3*σ*_c_. Then, the probability of survival for individual *i* is 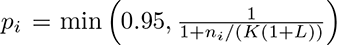, where *K* and *L* are parameters that are approximately equal to the the carrying capacity per unit area and the average lifetime at equilibrium, respectively. In our simulations *L* = 4 and the number of offspring per mating is Poisson distributed with mean 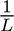. Edge effects are reduced, but not entirely avoided, by decreasing individual fitness proportional to the square root of distance from the habitat edges in units of *σ_c_.* When the proposed location of an offspring would have them disperse outside the bounds of the habitat, the individual is not born, and they are not replaced by another random location. We used a genome length of 10^8^ bp and recombination rate 10^-8^ crossovers per bp. This model was implemented in SLiM 3.7 (Haller and Messer, 2019). Model parameters varied between experiments and the relevant parameter ranges are described in Table 3.

**Table 3:** Parameter distributions used for simulation. The “Params.” column lists the identifier for the parameter set, which is referenced in the main text. “Description” is a brief description of the parameter set. “*σ*” is the distribution of the dispersal parameter. *“K”* is the major determinant of population density. *N*_1_ is the range of population sizes observed in the final (sampled) generation for each set of simulations. “Δ*_N_*” describes the history of population size change: for rows with braces, a random multiplier was chosen from one of two uniform distributions, each with probability 0.5. The ancestral *N_e_* was set to the multiplier x present day *N.* “Habitat width” is for the full habitat. “Samp. width” is the width of the sampling area as a proportion of the full habitat width. “Edge” is a distance from each side of the habitat that was excluded from sampling to avoid edge effects.

| Params. | Description | $\sigma$ | $K$ | $N_1$ | $\Delta N$ | Habitat width | Samp. width | Edge |
| --- | --- | --- | --- | --- | --- | --- | --- | --- |
| 1 | Comparing estimators | U(0.245, 3.67) | 5 | (10565, 12463) | constant | 50 | 1 | 3 |
| 2 | Baseline | U(0.245, 3.67) | 5 | (10565, 12463) | constant | 50 | 1 | 3 |
| 3 | Variable density | U(0.245, 3.67) | log-U(0.1, 20.0) | (87, 49200) | constant | 50 | 1 | 3 |
| 4 | Large density | U(0.245, 3.67) | U(20.0, 40.0) | (45152, 97497) | constant | 50 | 1 | 3 |
| 5 | Demographic history | U(0.245, 3.67) | 5 | (10565, 12463) | $\begin{cases} U(\frac{1}{5}, 1) \\ U(1, 5) \end{cases}$ | 50 | 1 | 3 |
| 6 | Extreme $N$ change | U(0.245, 3.67) | 5 | (10565, 12463) | $\begin{cases} U(\frac{1}{10}, \frac{1}{5}) \\ U(5, 10) \end{cases}$ | 50 | 1 | 3 |
| 7 | Variable habitat size | U(0.245, 3.67) | 2 | (245, 44695) | constant | U(15, 150) | 1 | $\sigma$ |
| 8 | Large habitat size | U(0.245, 3.67) | 2 | (6030, 176023) | constant | U(150, 300) | 1 | $\sigma$ |
| 9 | Variable sampling width | U(0.245, 3.67) | 5 | (10565, 12463) | constant | 50 | U(0.2, 0.8) | $\sigma$ |
| 10 | Large sampling width | U(0.245, 3.67) | 5 | (10565, 12463) | constant | 50 | U(0.8, 1.0) | $\sigma$ |
| 11 | Multiple nuisance par. | log-U($1.2 \times 10^{-3}$, $1.2 \times 10^2$) | log-U($10^{-3}$, $10^4$) | (163, 301891) | $\begin{cases} U(\frac{1}{5}, 1) \\ U(1, 5) \end{cases}$ | log-U(2, $10^3$) | U(0.0, 1.0) | $\sigma$ |

In the current paper, we aim to compare `disperseNN` directly with the methods from Rousset (1997) and Ringbauer et al. (2017), who estimate *effective* dispersal. Effective dispersal describes the movement of genetic material over generations and is usually measured as the root mean squared parent-offspring directional displacement, *σ* (i.e., using a model where the displacement in any particular direction has standard deviation *σ,* and the root mean squared Euclidean distance between parent and offspring is 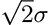). In our forward-in-time simulations, *σ* depends on each of the previously mentioned individual-based processes to some extent, which makes it an outcome of the simulation instead of a model parameter. The parameters *σ_f_* and *σ_m_* are the primary determinants of *σ,* while the competitive interaction distance, *σ_c_,* has a much smaller effect on *σ*: varying *σ_c_* tenfold changes *σ* less than 1% (Figure S9). Since *σ* is influenced by more than one parameter, we set *σ_f_ = σ_m_ = σ_c_* in the training simulations for `disperseNN`, which results in a simpler model and a relatively uniform distribution for *σ*.

In the experiments we present in the main text, dispersal history was not recorded during simulations. Therefore, we provide *σ_f_ as* the target to `disperseNN` during training, and apply a post hoc correction to rescale the predicted values: 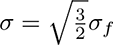. This correction is appropriate because the mean squared directional displacement between mother and child is 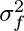, and between father and child is 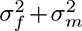. So, the mean squared directional displacement between a child and a randomly chosen parent is the average of these two, which if *σ_f_ = σ_m_* is equal to 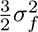. The corrected values are within a few percent of effective *σ* (Figure S10), allowing us to compare the `disperseNN` output directly with other genetics-based methods. This approach makes the assumption that each of the three types of local interactions occur on the same order, which seems reasonable for some species, but may be inappropriate for others. In a scenario where `disperseNN` is trained with *σ_f_= σ_m_* = *σ_c_*, but this assumption is not met in the test data, the *σ* estimate with post hoc correction still works but may be moderately biased (Figure S11).

For Supplementary Figures S9, S10, and S11 we tracked effective dispersal during the simulation as follows: let *f_i_* be the total number of offspring of individual *í* (acting as either father or mother) and 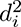 be the average squared displacement along an axis between individual *i* and their parent, averaged over parents and choice of axis (*x* or *y*). We then estimated the effective dispersal distance as 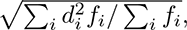 where the sum is over all individuals alive at any point in the simulation. In experiments where one or more of {*σ_f_, σ_m_, σ_c_*} were varied independently of the others, each parameter was drawn from a Uniform(0.2, 3).

After the completion of the spatial, forward-in-time SLiM simulation, initial genetic diversity was produced using a coalescent simulation in msprime (Haller et al., 2018). This strategy, known as “recapitation”, was necessary to reduce computation time to manageable levels, as the coalescent stage of the simulation is much faster than the spatially-explicit portion. The ancestral *N_e_* was set to the “present day” census population size for recapitation, as this quantity is more easily observable than effective population size. This portion of the simulation proceeded until all genealogical trees had coalesced. Thus, the complete simulation involves random mating for older generations equivalent to a Wright-Fisher model, with a number of recent generations that are spatially explicit (Table 4). Most of our experiments used 100,000 spatial-SLiM generations. However, to facilitate larger simulations for the multiple nuisance parameters experiment (Parameter Set 11), we ran only 100 generations of spatial SLiM due to computational limitations. This may seem too few, however since spatial mixing happens over shorter timescales than that of coalescence, then it seems likely that the signal that we are interested in for estimating dispersal would be generated over the recent past. We verified that `disperseNN` can predict *σ* from full-spatial test data after training on simulations with only 100 spatial generations, although *σ* was moderately underestimated (Figure S12). To simulate population size changes, we recapitated with msprime as before, but included an instantaneous decline or expansion between 100 and 100,000 generations in the past.

**Table 4:** Analysis parameters. The “Params.” column lists the identifier for the parameter set, which is referenced in the main text. “Description” is a brief description of the parameter set. “Sims.” is the number of true replicates, i.e., SLiM simulations, represented in training. “Training” is the size of the total training set after drawing multiple samples from each simulation. “Spatial gen.” is the number of spatial generations simulated in SLiM. “n” is the sample size. “SNPs” is the number of SNPs used in training. “Phased” describes whether the data were phased or not for training. See Table 3 for corresponding distributions for parameters of the simulation model.

| Params. | Description | Sims. | Training | Spatial gen. | $n$ | SNPs | Phased |
| --- | --- | --- | --- | --- | --- | --- | --- |
| 1 | Comparing estimators | 1,000 | 50,000 | 100,000 | 10 and 100 | $2.5 \times 10^5, 5 \times 10^5$ | Y |
| 2 | Baseline | 1,000 | 50,000 | 100,000 | 100 | 5,000 | Y |
| 3 | Variable density | 1,000 | 50,000 | 100,000 | 100 | 5,000 | Y |
| 4 | Large density | 1,000 | 50,000 | 100,000 | 100 | 5,000 | Y |
| 5 | Demographic history | 1,000 | 50,000 | 1,000 | 100 | 5,000 | Y |
| 6 | Extreme $N$ change | 1,000 | 50,000 | 1,000 | 100 | 5,000 | Y |
| 7 | Variable habitat size | 1,000 | 50,000 | 100,000 | 50 | 5,000 | Y |
| 8 | Large habitat size | 1,000 | 50,000 | 100,000 | 50 | 5,000 | Y |
| 9 | Variable sampling width | 1,000 | 50,000 | 100,000 | 100 | 5,000 | Y |
| 10 | Large sampling width | 1,000 | 50,000 | 100,000 | 100 | 5,000 | Y |
| 11 | Multiple nuisance par. | 2,300 | 100,000 | 100 | U-int(10,100) | 5,000 | N |

Individuals were sampled randomly from within the sampling window. When the specified size of the spatial sampling window was smaller than the full habitat, the position of the sampling window was chosen randomly, with *x* and *y* each distributed uniformly (Figure 5), excluding edges. The amount of edge cropped was either set to (i) *σ* for each simulation, or (ii) the maximum of the simulated *σ* range for the whole training set, depending on which simulation parameters were free to vary; the latter was necessary to avoid information leakage during training. For some analyses, multiple, partially-overlapping samples were drawn from the same simulation to save computation time; these cases are noted in Table 4. This strategy allows for large training sets to be generated from a smaller number of starting simulations.

**Figure 5:**
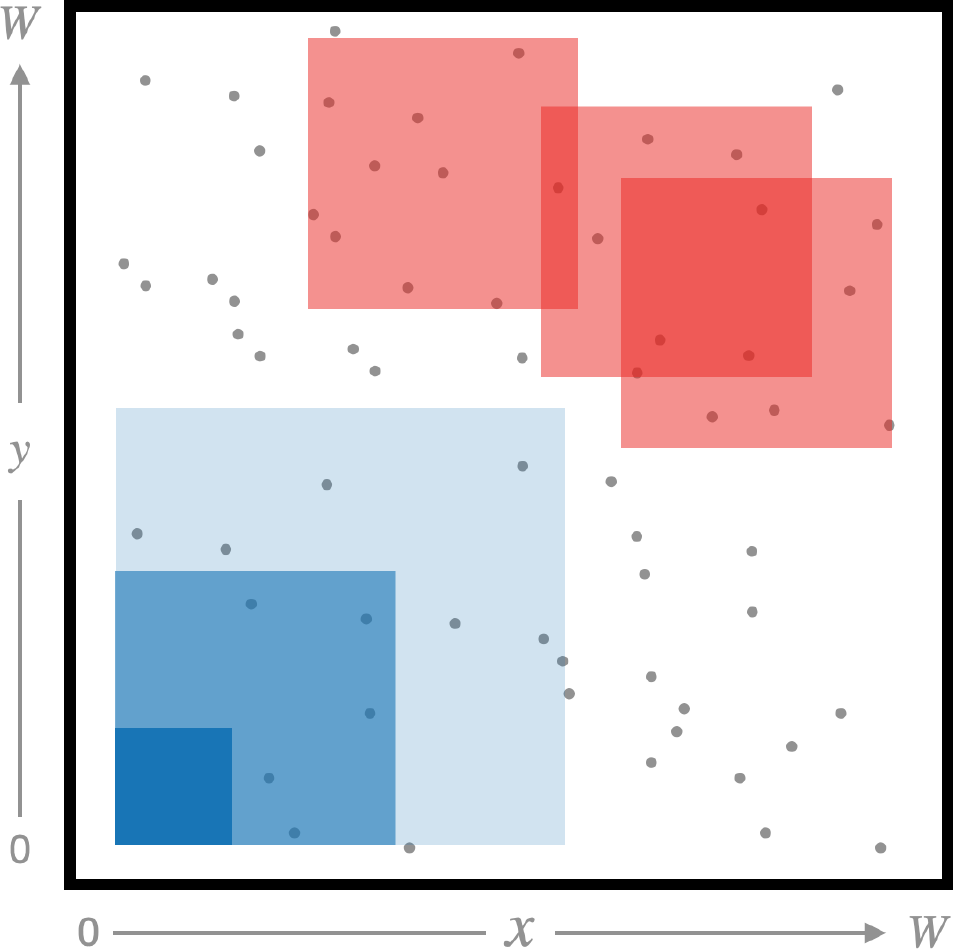
Cartoon showing different sampling strategies. The black box represents the full simulated habitat. For some experiments, we both (i) varied the width of the square sampling window—blue boxes show examples of differing sampling widths—, and (ii) assigned a uniform-random position for the sampling window—red boxes show different positions for the sampling window.

To obtain genetic data at *m* varying sites, neutral mutations were superimposed on the tree sequences using msprime vl.0 (Baumdicker et al., 2022) (for values of *m* see the “SNPs” column in Table 4). This mutate-afterwords approach is efficient because we only add mutations that affect the sampled individuals (Kelleher and Lohse, 2020). For efficiency we used an iterative approach for adding mutations because our analyses used a fixed, and usually small, number of mutations. Specifically, we started by simulating mutations with a very small mutation rate, 10^-15^. If we did not yet have *m* SNPs, we increased the mutation rate by 10x, and added additional mutations with the updated mutation rate. This was iterated until at least *m* mutations had been obtained. Finally, a uniform sample of *m* biallelic SNPs were sampled to represent the genotype matrix input to `disperseNN`. The result of this procedure is that the genotype matrix for each simulated dataset contains the same number of SNPs, *m,* regardless of the actual number of variable sites in the sampled individuals, and irrespective of mutation rate. Thanks to the Poisson nature of neutral mutations, this procedure is equivalent to having simulated with a higher mutation rate and randomly selected *m* variable sites.

The input for `disperseNN` consists of two things: the width of the spatial sampling area, and a genotype matrix, having one row for each SNP and one or two columns per individual depending on the phasing designation. If phased, the genotype matrix contained two columns per individual, randomly ordered, with Os and Is encoding minor and major alleles, respectively. If unphased, the genotype matrix contained one column per individual with genotypes encoded as Os, Is, and 2s, representing the count of the minor allele. In order to facilitate various sample sizes in real applications, our pre-trained model used a random sample size during training, 10 ≤ n ≤ 100, with zero padding out to 100 columns. To obtain the second input, we used the furthest distance between pairs of samples as the sampling width. The training targets are the true values of log(*σ*). Thus, the output from the CNN is in log space (`disperseNN` exponentiates the result before writing the predictions).

In generating training data for the pre-trained network, we sought to explore a large parameter range: each parameter varied over several orders of magnitude (Parameter Set 11). However, swaths of parameter space described by the ranges in Table 3 were not represented in the training data, due to the following logistical hurdles. First, simulations where the population died were not included in the training set. The excluded simulations had small carrying capacity and small habitat size, or small habitat size and large *σ*, for example. Next, some simulations could not be run due to computational constraints: maximum RAM of 175 gigabytes and two-week wall time on our computing cluster. For example, combinations of large carrying capacity and large habitat size were not simulated. As a result, only 12% of attempted simulations were included in training, and for each parameter the *realized* distribution—representing successful simulations— differed from the distribution from which the model settings were drawn (Figure S6), which had been uniform in log space.

### CNN architecture and training

Tensorflow (Abadi et al., 2016) and Keras (https://github.com/keras-team/keras) libraries were used to develop `disperseNN`. The first input tensor, the genotype matrix, goes through successive convolution and pooling layers, a strategy that is characteristic of CNNs (Figure 1). We adjusted the number of convolution and pooling layers based on the size of the genotype matrix: the number of convolution layers assigned was equal to floor (log_10_ (number of SNPs))–1. The filter size of successive convolution layers was 64 for the first layer, and 44 larger for each successive layer. The convolution layers are one-dimensional, such that the convolution kernel spans all individuals (columns) and two SNPs (rows), with stride size equal to one. Average pooling layers were also one dimensional, spanning all individuals and 10 SNPs. After the convolutional portion of the network, the intermediate tensor was flattened and put through three fully connected layers each with 128 units and rectified linear unit (ReLU) activation. A second input branch was used for the sampling area. This input tensor with size = 1 was concatenated with the preceding branch, then subjected to a 128-unit dense layer with ReLu. Finally, a dense layer with linear activation was applied which outputs a single value, the estimate for *σ*.

During training we held out 20% of the training set for computing a validation-loss between epochs. We used a batch size of 40, mean squared error loss, and the Adam optimizer. The learning rate was initialized as 10^-3^. The “patience” hyperparameter determines both the length of training, and learning rate adaptations during training: after a number of epochs equal to patience/10 without improvement in validation loss the learning rate is halved, and training proceeds until a number of epochs equal to patience pass without improvement in validation loss. Patience was set to 100 for all training runs excluding the pre-trained model. For the pre-trained model, we explored a grid of different hyperparameter settings: patience values of 10, 20, 30, 40, and 50; initial learning rates of l0^-4^, 10^-3^, and 10^-2;^ and dropout proportions of 0, 0.1, 0.2, and 0.3. We landed on settings that consistently gave the lowest MRAE: patience = 10, initial learning rate of 10 ^3^, and 0 dropout.

The architecture we present does not use spatial information, except for a scalar for the width of the sampling area. Although we focus on the non-spatial architecture, an extension to our model might incorporate the relative spatial distances between samples. For the goal of conveying spatial information we have explored basic architectures that did not work well, at least with uniform sampling (see Appendix); however, the spatially-explicit architectures may help differentiate spatial sampling strategies (e.g., point sampling, transect sampling, etc.), which we have not yet explored. Many alternative architectures are conceivable and may be analyzed in future studies. For example, the spatial connectedness between individuals might be shown to the network by sorting the genotypes by their geographic coordinates.

### Comparison with other estimators

The method from Rousset (1997) uses the observation that under certain assumptions 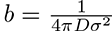, where *D* is density and *b* is the slope of the least squares fit of a*_r_*/(l – *a_r_*) to log geographic distance. Here, *a_r_* is a measure of genetic differentiation between two individuals (analogous to *F_st_*) that can be estimated for a pair of individuals, 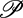, as 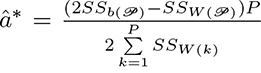. In this equation 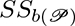 is the sum of squared differences between the two individuals’ genotypes, 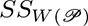 is the sum of squared differences between genomes within the individuals, *P* is the total number of pairs of individuals in the sample, and 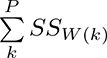 are within individual differences summed over the *P* different pairs of individuals. We applied Rousseťs method to the same genotypes and sample locations as for `disperseNN`. The value for *D* used with this method was calculated using the following procedure. First, *N_e_* was calculated using the “inbreeding effective size” from Waples (2002):

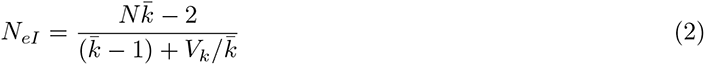

where 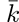 is the mean number of offspring per individual and *V_k_* is the variance in offspring number among individuals. To deal with overlapping generations, we calculated *N_eI_* each simulation cycle using census *N* from the current cycle along with the lifetime offspring count, *k,* for dying individuals. The per generation mean *N_eI_* was calculated at the end of the simulation. With an estimate for *N_e_* in hand, we calculate density *as D = N_e_*/(*W –* 2*σ*)^2^, where the 2*σ* is to exclude edges which have reduced density.

A second comparison was made with `IBD-Analysis` (Ringbauer et al., 2017). The authors used the distribution of identity-by-descent tract lengths shared between individuals to estimate *σ*. They derived analytical formulas describing the distribution of identity-by-descent tract lengths in continuous space and provided an inference scheme that uses maximum likelihood to fit these formulas. For our comparison we used two different analysis pipelines for `IBD-Analysis`. First, we extracted the true identity-by-descent tracts directly from the tree sequences output from SLiM. Specifically, for each pair of individuals, for each combination of chromosomes between the individuals, we simplified the tree sequence to represent only the recombination history between the two chromosomes, and extracted segments that were inherited from a common ancestor without recombination. These were the identity-by-descent tracts used as input for the `IBD-Analysis` program, which was obtained from https://git.ist.ac.at/harald.ringbauer/IBD-Analysis. Separately, we inferred identity-by-descent tracts in the simulated data using an empirical tool, `Refined IBD` (Browning and Browning, 2013), and used the inferred identity-by-descent blocks as input `IBD-Analysis`. We used the following `IBD-Analysis` settings: genome length of le8 bp, minimum block length of 1 cM (default). We used the following `Refined IBD` settings: minimum block length of 1 cM (default=1.5 cM).

### Empirical data

To demonstrate the utility of `disperseNN`, we applied it to preexisting publicly available datasets that have the following criteria: spatially distributed genetic data, latitude and longitude metadata available, ten or more sampling locations, sampling area less than 1000 km, at least 5,000 biallelic SNPs, and a ready-to-plug-in SNP table that had been processed and filtered by the original authors. For some datasets with overall sampling width more than 1000 km, we were able to subset to a smaller cluster of sample locations (see details specific to each dataset below). When multiple individuals were sampled from the same location we chose one random individual from each location, in order to better match the sampling scheme used in generating training data. SNP tables were converted to genotype matrices after minimal processing: we removed indels and sites with only one, or more than two, alleles represented in the sampled subset. We required all sampled individuals to be genotyped to retain a SNP, except when we note otherwise—see details specific to each dataset below.

Mosquito data were downloaded following instructions from https://malariagen.github.io/vector-data/ag3/download.html. We used a dense cluster of sampling localities in Cameroon that had been identified as *Anopheles gambiae.* Individual VCFs were merged using `bcftools` (vl.14). Chromosomes 3L and 3R were analyzed; 2L and 2R were excluded due to previously reported large inversions (Lobo et al., 2010; Riehle et al., 2017).

*Arabidopsis* data were downloaded from https://100lgenomes.org/data/GMI-MPI/releases/v3.1/ as a single VCF. Two conspicuous geographic clusters were chosen from Sweden and Spain to minimize the geographic sampling area. All five chromosomes were analyzed.

Sunflower data were downloaded following instructions from https://rieseberglab.github.io/ubc-sunflower-genome/documentation/. Geographic clusters of sampling localities were identified in Texas (*Helianthus argophyllus*) and on the border of Kansas and Oklahoma (*H. petiolaris*). Individual VCFs were merged into multi-sample VCFs for each of the two species. Chromosomes 1-17 were analyzed, excluding a number of unplaced scaffolds.

VCFs for oyster (*Crassostrea virginica*; Bernatchez et al. (2019)), bumble bee (*Bombus*; Jackson et al. (2018)), Atlantic halibut (*Hippoglossus hippoglossus;* Kess et al. (2021)), white-footed mouse (*Peromyscus leucopus;* Munshi-South et al. (2016)), Réunion grey white-eye (*Zosterops borbonicus*) and Réunion olive white-eye (*Zosterops oliυaceus;* Gabrielli et al. (2020)), and wolf (*Canis lupus;* Schweizer et al. (2016)) were downloaded directly from The Dryad Digital Repository. Clusters of sample locations were chosen in each dataset to maximize sampling density. In the datasets from *Bombus υosnesenskii, Peromyscus leucopus, Zosterops borbonicus,* and *Zosterops oliυaceus,* we allowed as few as 85%, 60%, 90%, and 90% of individuals to be genotyped to retain a SNP, respectively, and missing genotypes were filled in with the major variant.

To calculate the width of the sampling window for empirical data, we calculated the geodesic distance between each pair of individuals using the package `geopy` with the WGS84 ellipsoid. This distance represents the shortest path on the surface of the Earth between points. The longest distance between pairs of sample locations was used as the sampling width, which we provided in kilometers to `disperseNN`.

## Acknowledgements

We thank John Novembre and anonymous reviewers for their constructive feedback, in particular their ideas for leveraging spatial information. We are thankful to Harold Ringbauer for assistance with the methods comparison analysis. And we thank members of the Kern-Ralph Co-lab, as well as Dan Schrider, Will Booker, and Ryan Gutenkunst for valuable input along the way. Computation was done using the University of Oregon’s cluster, Talapas, with help from the Research Advanced Computing Services team. This work was supported by the National Institutes of Health [grant numbers F32GM146484 to C.S. and R01HG010774 to A.K.].

## Data availability

The `disperseNN` code is available on GitHub at the following link: https://github.com/kr-colab/disperseNN.

## Supplementary material

**Figure S1:**
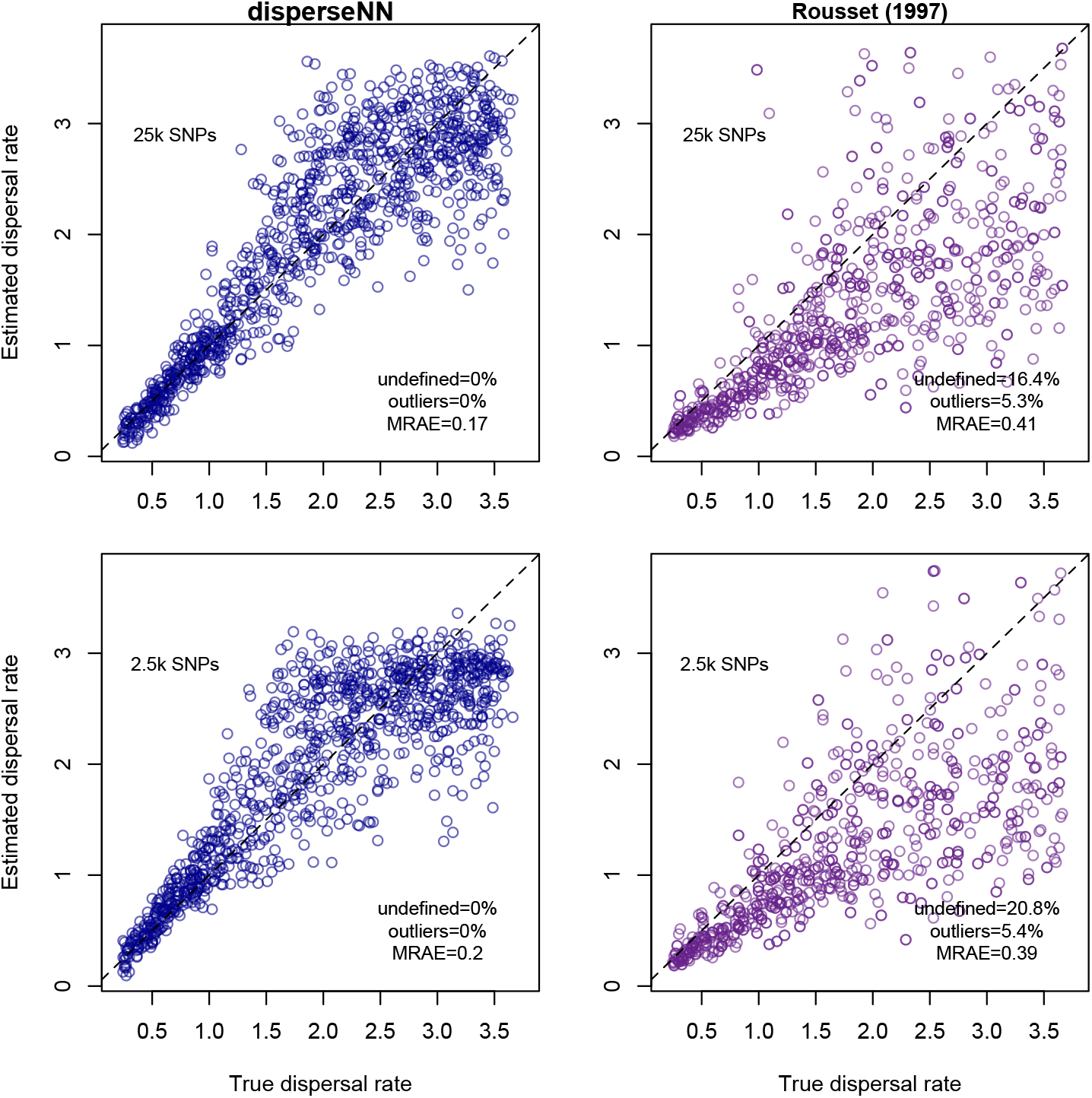
Comparison between `disperseNN` and Rousseťs method using 10 individuals and varying SNP number (other parameters as in Parameter Set 1); compare to Figure 2. MRAE is the mean relative absolute error.

**Figure S2:**
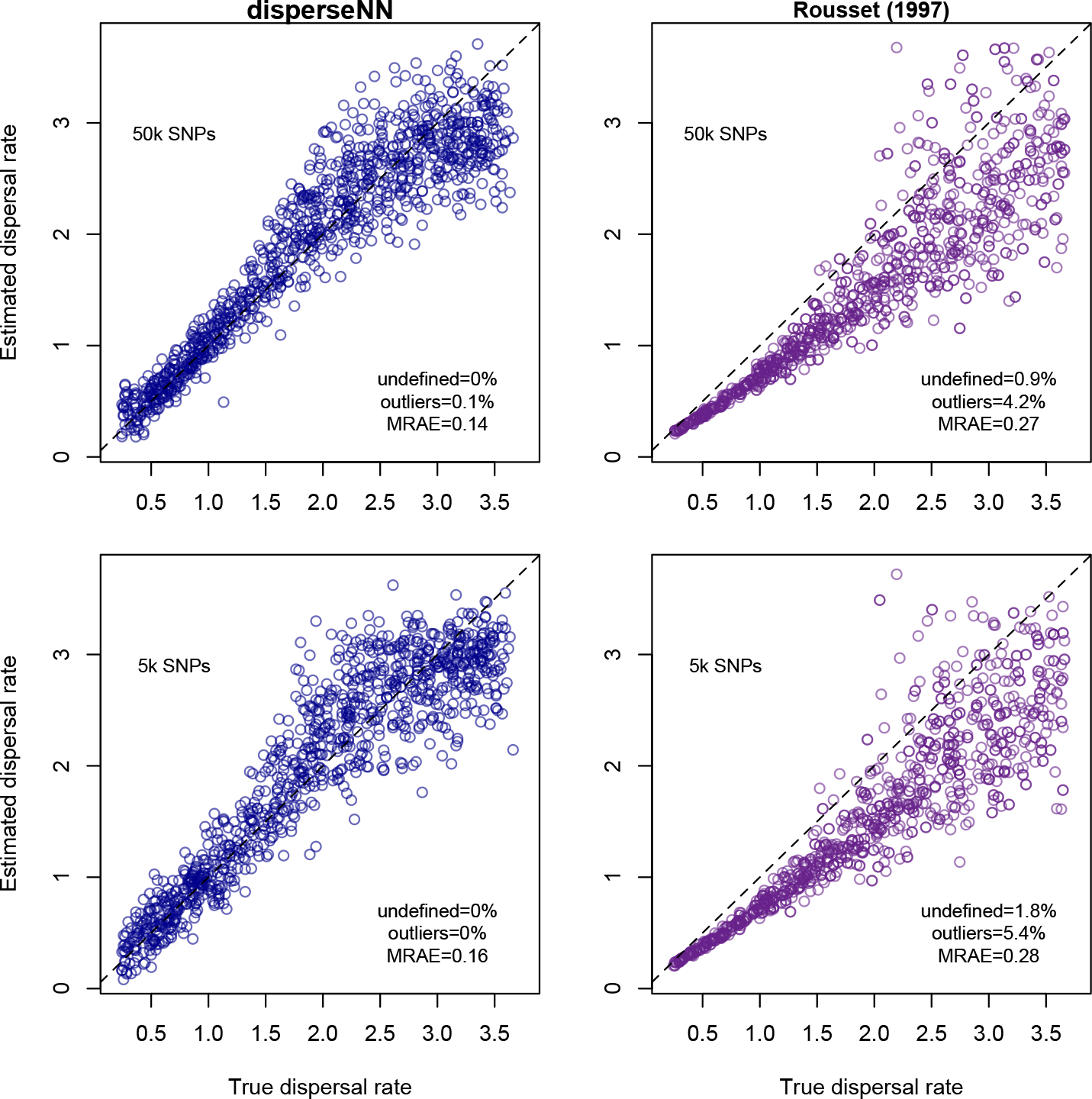
Comparison between `disperseNN` and Rousseťs method using 100 individuals and varying SNP number (other parameters as in Parameter Set 1); compare to Figure 2. MRAE is the mean relative absolute error.

**Figure S3:**
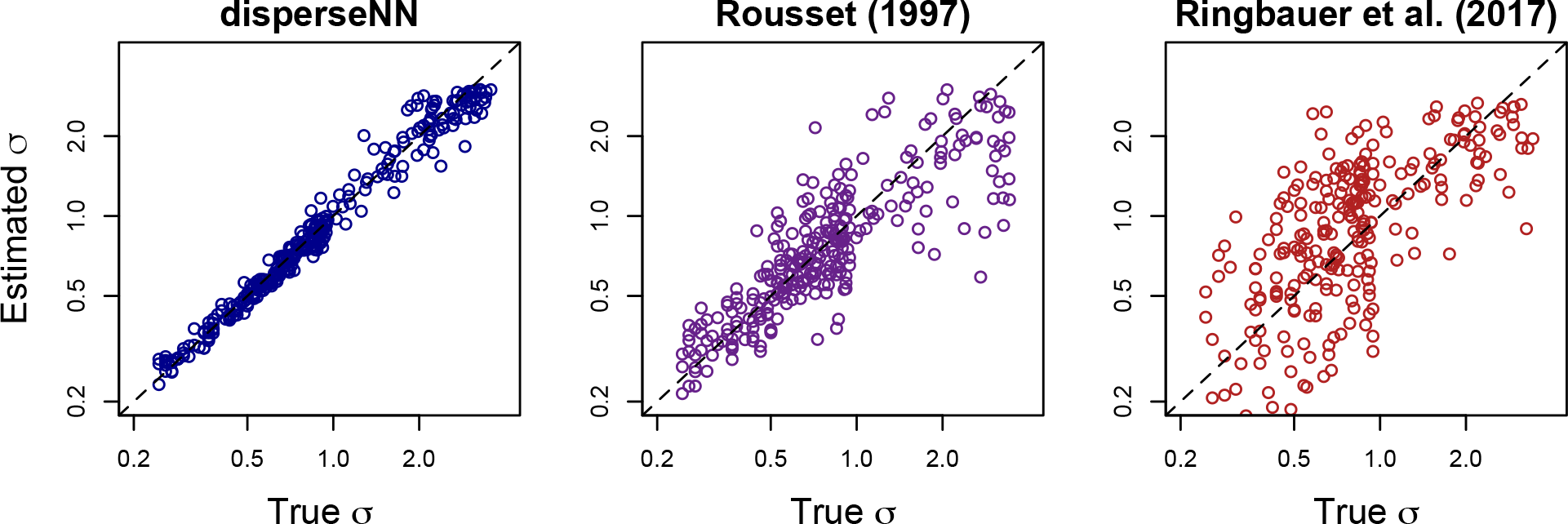
Predictions with log-transformation to show relative error (*n =* 10; Parameter Set 1). Data points in the larger half of the log(*σ*) range were down-sampled to one-half the number of points in the smaller half of the range to obtain roughly even density of points across the full range; before down-sampling, points were more dense towards the right-hand side which might give the false impression of larger variance.

**Figure S4:**
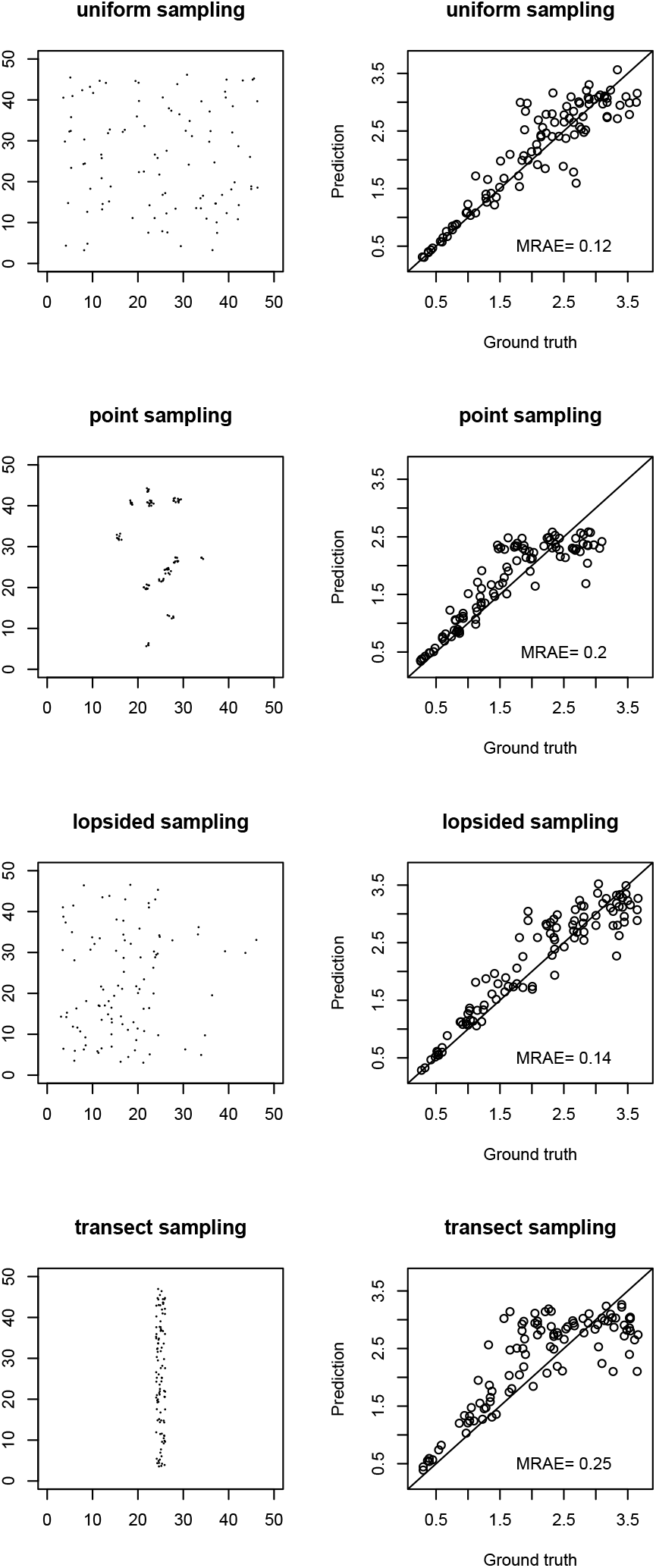
Exploring different sampling strategies. The left column shows the geographic distribution of sampling locations and the right column shows the `disperseNN` output, having trained with Parameter Set 2. For the ‘point sampling’ scenario, we used the sampling configuration from Munshi-South et al. (2016); for this, we used the same subset of 12 sample locations used in our empirical analysis, except here we used more than one individual per sampling locality, ranging from n = 4 to n = 11 per location. MRAE is the mean relative absolute error.

**Figure S5:**
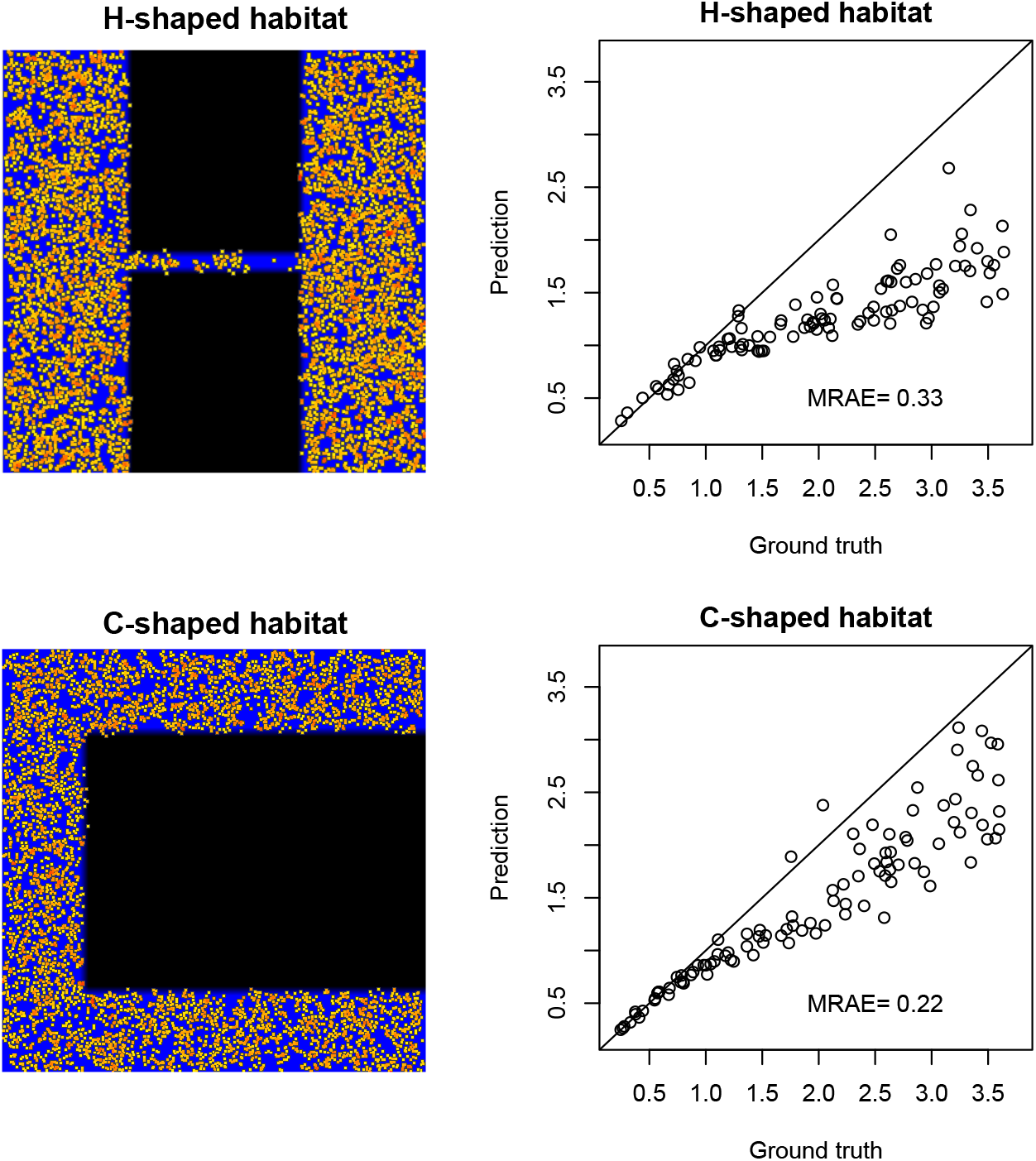
Exploring misspecified habitat shapes. The left column shows the simulation that produced the test data (screenshots of SLiM GUI): a square map (50 units wide) where individuals are yellow/orange, suitable habitat is blue, and unsuitable habitat is black. The right column shows the `disperseNN` output, having trained with a uniform (all blue) map (Parameter Set 2). The H-shaped habitat was inspired by McRae (2006) and the C-shaped habitat was inspired by McRae and Beier (2007). MRAE is the mean relative absolute error, and both axes are in units of *σ*.

**Figure S6:**
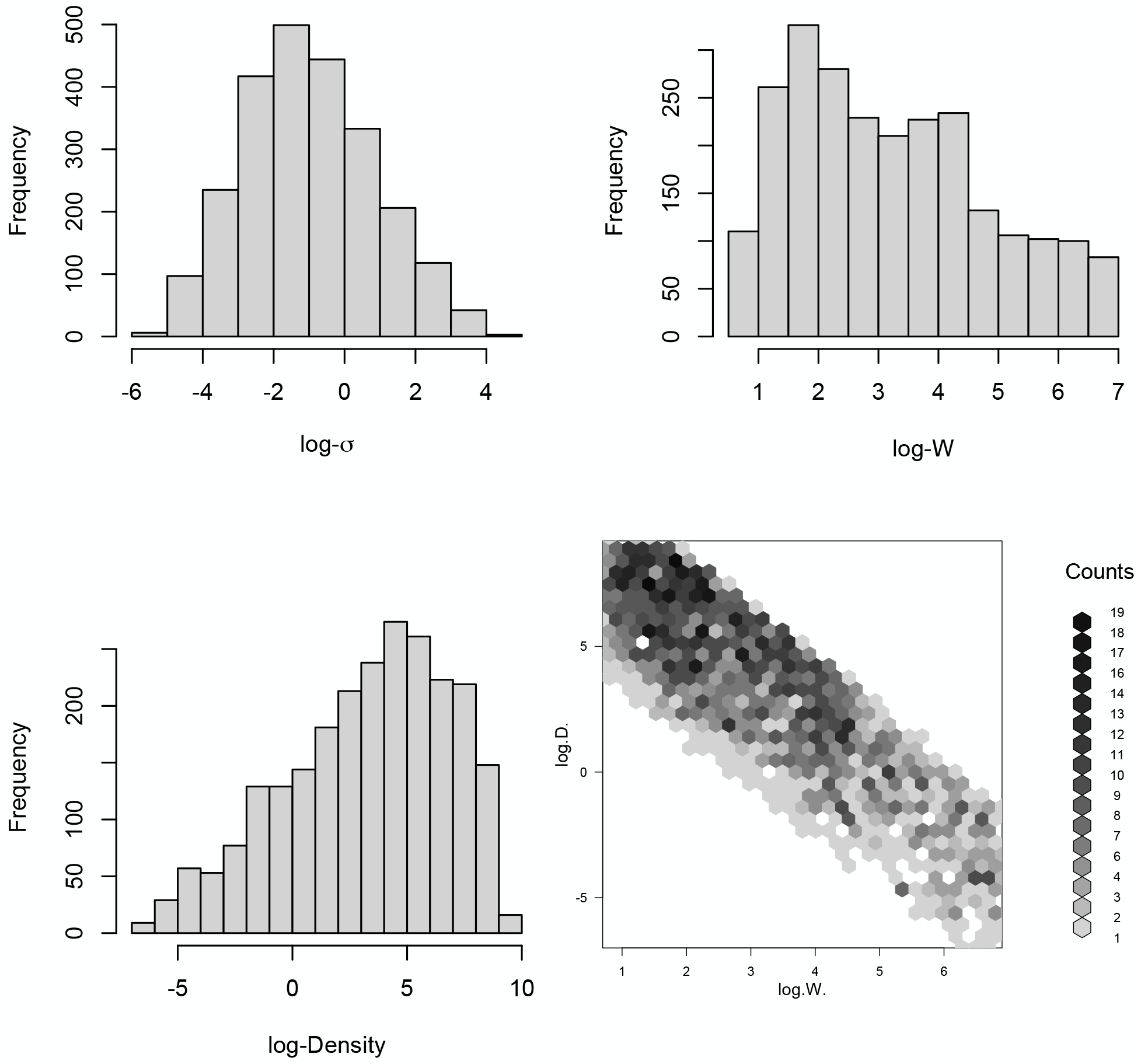
Realized training distributions for empirical analysis (Parameter Set 11). “D” is population density. “W” is habitat width. Some areas of parameter space could not be simulated due to population extinction or computational limitations.

**Figure S7:**
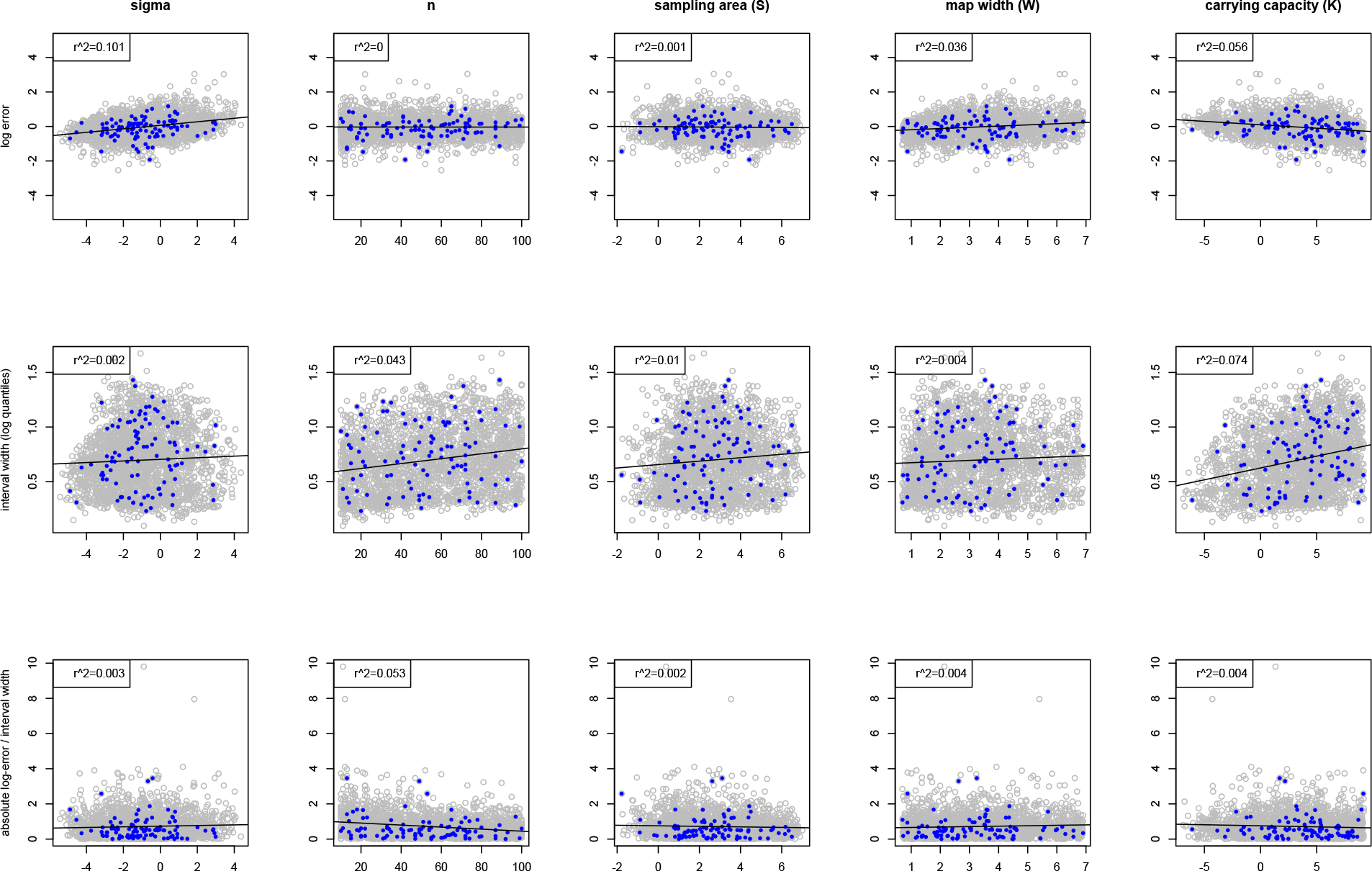
Exploring associations between five different predictor variables—(1) *σ*, (2) *n,* (3) sampling area, (4) map width, (5) carrying capacity—and three different response variables (A) log error, (B) the interval width of the log-transformed middle 95% range of the bootstrap distribution, and (C) absolute log-error divided by the interval width (Parameter Set 11). Shown are 2400 datasets including both held-out test data (blue; 100 datasets) and training data (grey; 2,300 datasets). The line of best fit and *r^2^* include all 2400 data points.

**Figure S8:**
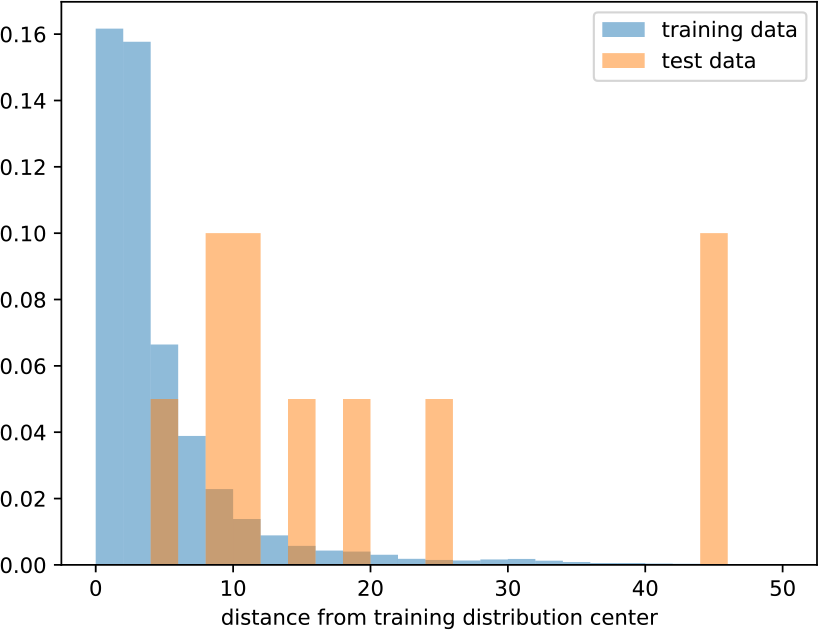
Mahalonobis distance from the center of the training distribution with respect to five summary statistics: nucleotide diversity, Tajima’s D, inbreeding coefficient, observed heterozygosity, and expected heterozygosity (Parameter Set 11). “test data” are the empirical datasets.

**Figure S9:**
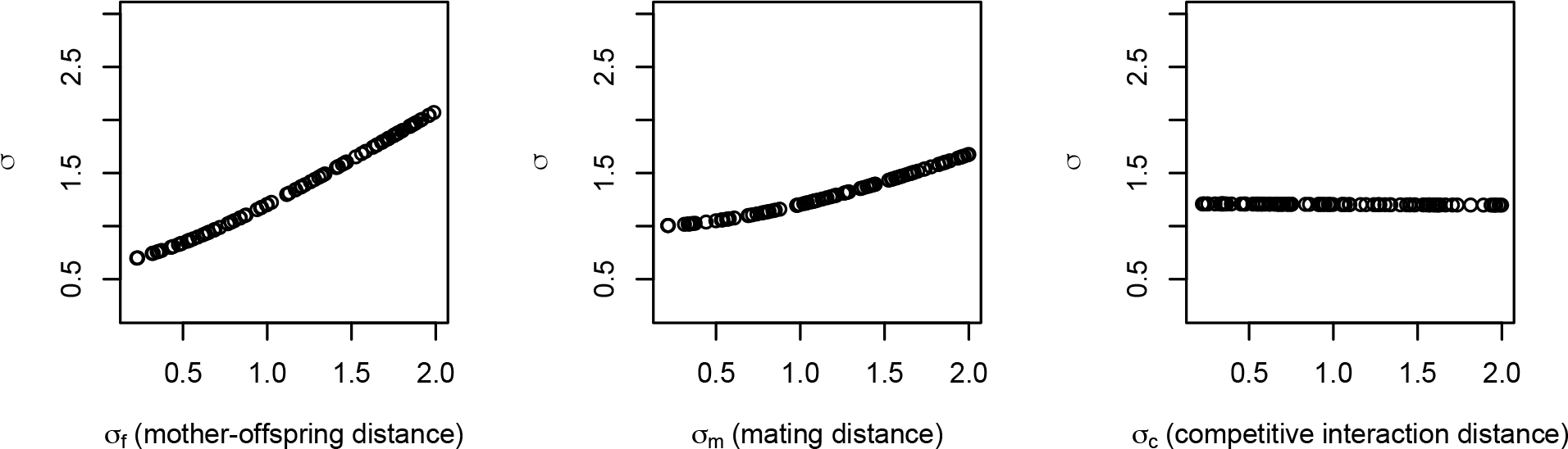
Visualizing true *σ,* while varying one of three spatial interaction distances at a time: mother­offspring distance (*σ_f_*), mating distance (*σ_m_*), and competitive interaction distance (*σ*_c_). Left plot: *σ_m_ = U*(0.2, 2); *σ_f_* = 1; *σ_c_* = 1. Middle plot: *σ_m_ =* 1*; σ_f_ = U*(0.2, 2); *σ_c_ =* 1. Right plot: *σ_m_ =* 1; *σ_f_ =* 1; *σ_c_* = *U*(0.2, 2).

**Figure S10:**
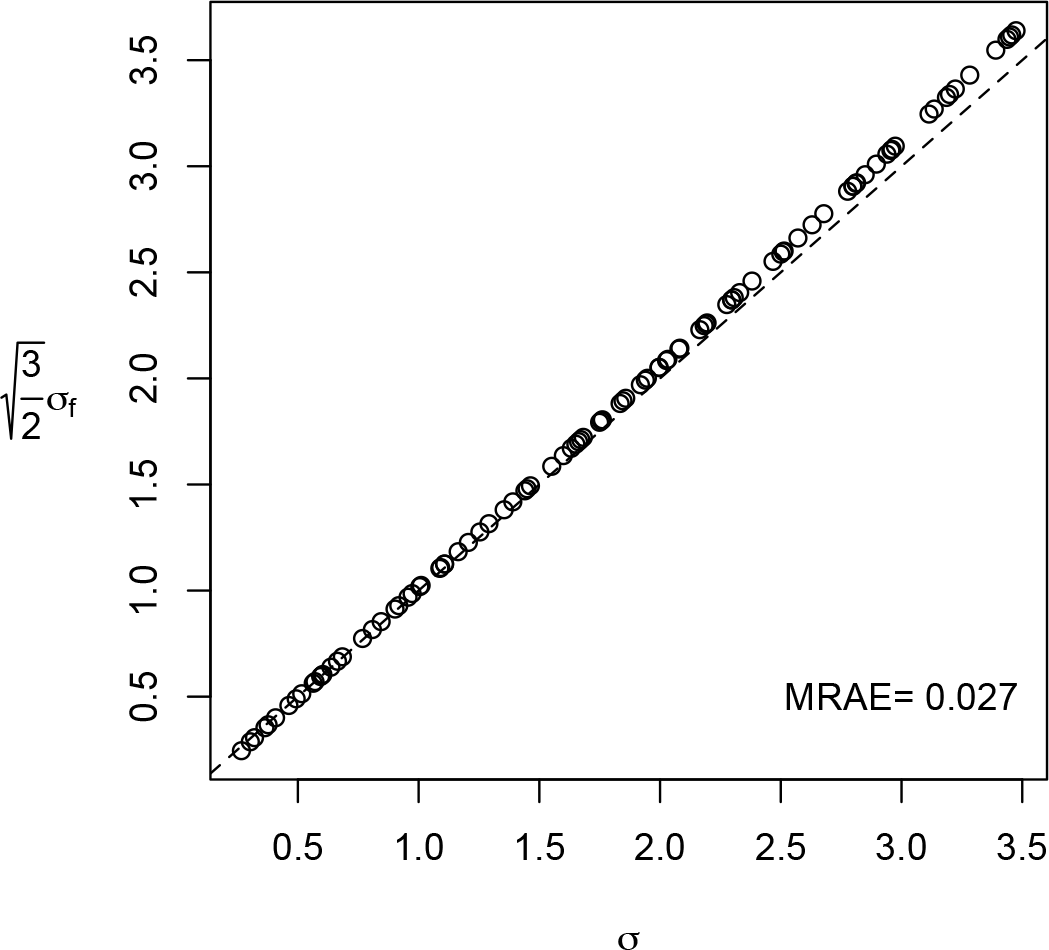
The x axis is true *σ* measured during the simulation, whereas the y axis shows true *σ_f_* with the post hoc correction. In these simulations *σ_f_ = σ_m_ = σ_c_* (mother-offspring distance *σ_f_*; mating distance *σ_m_*; competitive interaction distance *σ_c_*). MRAE is the mean relative absolute error.

**Figure S11:**
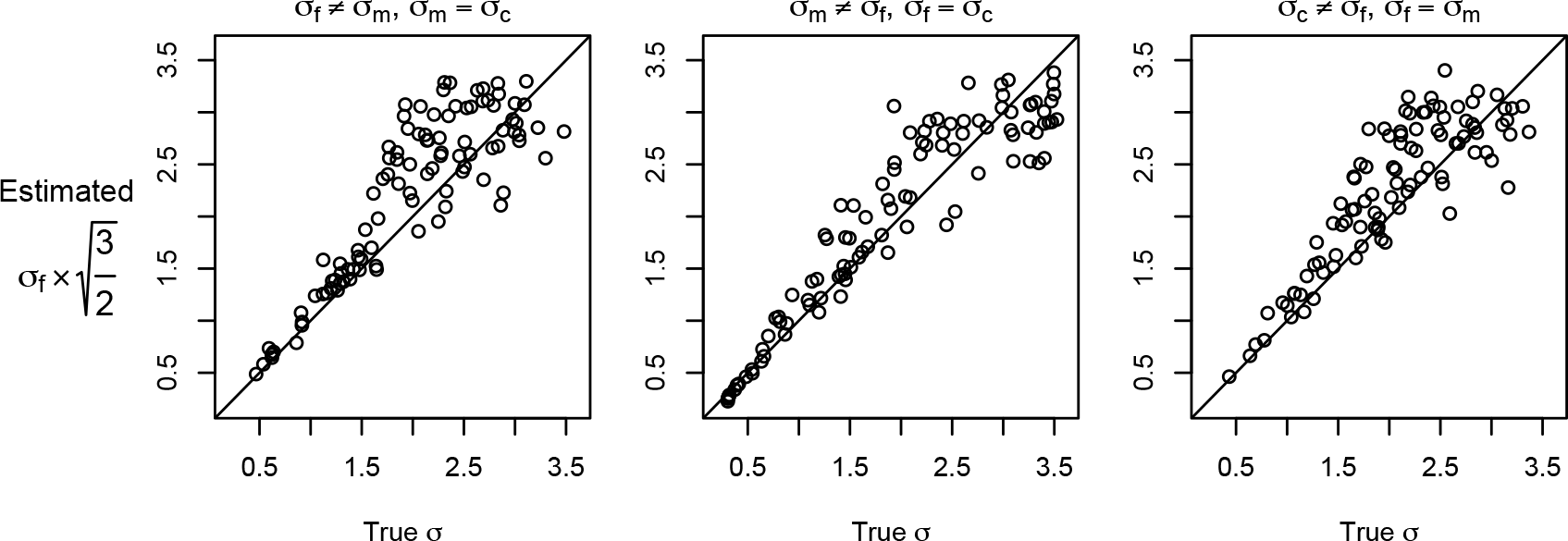
The x axis is true *σ,* and the y axis is estimated *σ_f_* multiplied by the post hoc correction, after training with *σ_f_ = σ_m_ = σ_c_* (mother-offspring distance *σ_f_*; mating distance *σ_m_*; competitive interaction distance *σ_c_*). In the test data for each panel, either *σ_f_, σ_m_,* or *σ_c_* was varied independently (Uniform(0.2,3)) of the other two spatial parameters which were set to be equal (drawn from another Uniform(0.2,3)).

**Figure S12:**
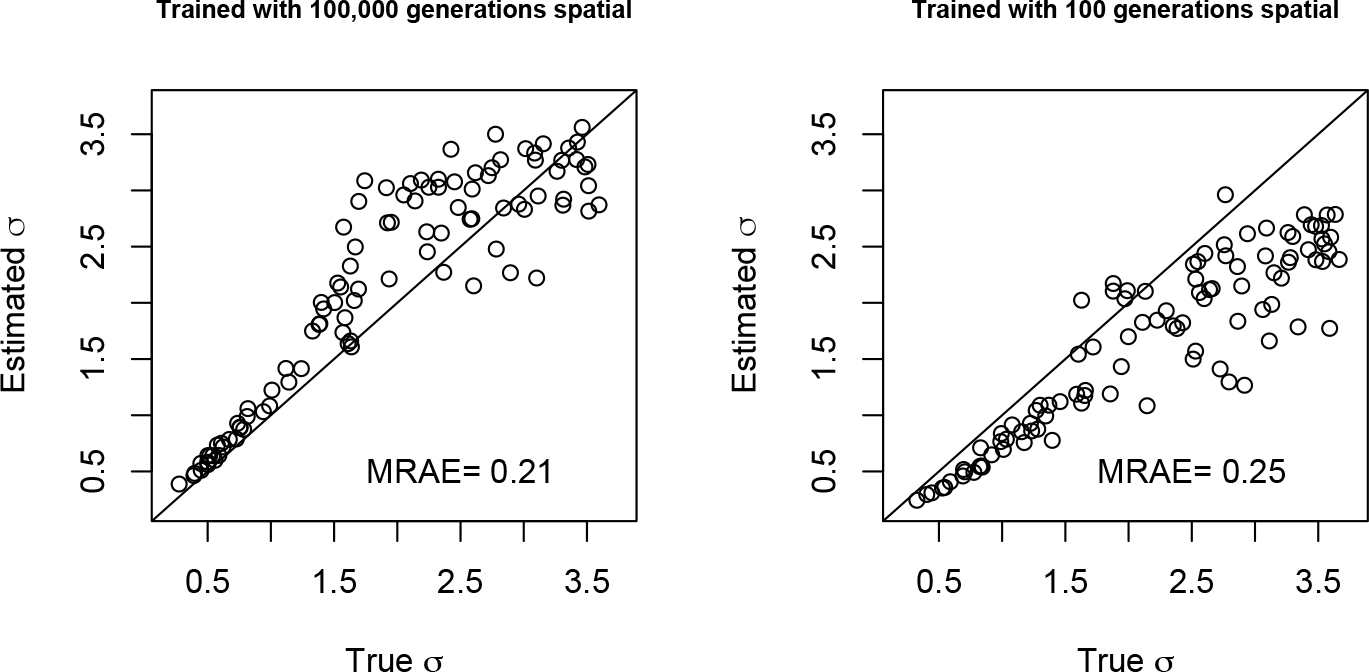
`disperseNN` trained with only 100,000 generations, or 100 generations, in spatial SLiM before recapitation with msprime (other parameters as in Parameter Set 2). Test data were full-spatial: SLiM was run until every tree had coalesced. MRAE is the mean relative absolute error.

## Appendix: Additionally investigated inputs to the method

This Appendix describes analyses not included in the main document, including strategies that did not work.

### A1. Attempts to use sampling localities

It is intuitive that signal about dispersal might be gleaned from the individual sample locations, as previous population-genetics-based inference methods use sample locations as input. We tried the following strategies for showing the sample locations to the CNN. In each experiment, we modified the neural network architecture to accommodate the sample locations in various ways. Otherwise, the neural network in each experiment closely resembled the architecture described in the main text.

- *Table of locations.* An *n x 2* array containing the *x* and *y* coordinates was shown to the CNN in a separate input branch (in place of the sampling width input). This input went through a single 128-unit dense layer with ReLu activation before flattening and concatenating with the previous branch.
- *Stored in genotype matrix.* Additional rows in the genotype matrix were used to store the *x* and *y* coordinates for each individual.
- *3-channel array.* A 3-dimensional array was used to store (1) the genotypes, (2) *x* coordinates, and (3) *y* coordinates. In the second and third channels, the spatial coordinates were repeated for *m* rows equal to the number of SNPs. Here, the neural network used ID-convolution and pooling layers, as described in the main text, however the convolution and pooling layers spanned all three channels simultaneously.
- *2D CNN.* We also tried a variation of the the 3-channel-array strategy using 2D-convolution and pooling layers with a 2×2 window.

For each of the above strategies, we trained the neural network in the same manner as the “baseline” model from the misspecification analysis in the main text. The outcome for each was the same: the mean RAE was indistinguishable from the baseline model that does not include sample locations. Moreover, we shuffled the sample locations input, such that each individual has a randomly assigned location, and the output was unchanged. Our interpretation is that the CNN ignores the location data in the experiments attempted thus far, either because the locations are not necessary for estimating *σ*, or because we failed to effectively show the network the locations.

### A2. Including isolation-by-distance summary statistics

We tested whether isolation by distance information in the form of summary statistics would improve inference of *σ*. Specifically, we summarized isolation-by-distance as:

- *b,* the slope of the line of best fit to genetic distances versus geographic distances.
- *r*^2^, the coefficient of correlation between genetic distance and geographic distance.

Including either (or both) of these statistics as a separate input branch of size one (or two) marginally improved validation accuracy. The new input branch went through a 128-unit dense layer with ReLu activation before concatenating with the previous branch. Thus, future empirical applications might explore using the above or different summary statistics alongside the genotype matrix for estimating *σ*, or other population genetic parameters. We did not present these results in the main text because (1) the benefit was negligible, and (2) it is beyond the scope of our study to decide on the most relevant and appropriate summary statistics, as countless other statistics might be evaluated for use with, or without, the genotype matrix that we used.

